# System-Level Quantification and Phenotyping of Early Embryonic Morphogenesis of *Caenorhabditis elegans*

**DOI:** 10.1101/776062

**Authors:** Guoye Guan, Ming-Kin Wong, Vincy Wing Sze Ho, Xiaomeng An, Lu-Yan Chan, Binghui Tian, Zhiyuan Li, Leihan Tang, Zhongying Zhao, Chao Tang

## Abstract

Cell lineage consists of cell division timing, cell migration and cell fate, and is highly conserved during development of nematode species. An outstanding question is how differentiated cells are genetically and physically regulated in order to migrate to their precise destination among individuals. Here, we first generated a reference embryo using time-lapse 3 dimensional images of 222 wild-type *C. elegans* embryos at about 1.5-minute interval. This was achieved by automatic tracing and quantitative analysis of cellular phenotypes from 4- to 24-cell stage, including cell cycle duration, division orientation and migration trajectory. We next characterized cell division timing and cell kinematic state, which suggests that eight groups of cells can be clustered based on invariant and distinct division sequence. Cells may still be moving while others start to divide, indicating strong robustness against motional noise in developing embryo. We then devised a system-level phenotyping method for detecting mutant defect in global growth rate, cell cycle duration, division orientation and cell arrangement. A total of 758 genes were selected for perturbation by RNA interference followed by automatic phenotyping, which suggests a cryptic genetic architecture coordinating early morphogenesis spatially and temporally. The high-quality wild-type reference supports a conceptual close-packing model for cell arrangement during 4- to 8-cell stage, implying fundamental mechanical laws regulating the topological structure of early *C. elegans* embryo. Also, we observed a series of remarkable morphogenesis phenomena such as induced defect or recovery from defect in mutant embryo. To facilitate use of this quantification system, we built a software named ***STAR 1.0*** for visualizing the wild-type reference and mutant phenotype. It also allows automatic phenotyping of new mutant embryo. Taken together, we not only provide a statistical wild-type reference with defined variability, but also shed light on both genetic and physical mechanisms coordinating early embryonic morphogenesis of *C. elegans*. The statistical reference permits a sensitive approach for mutant phenotype analysis, with which we phenotype a total of 1818 mutant embryos by depletion of 758 genes.

**Highlights & Graphical Abstract:**

● Spatial-Temporal Wild-Type Reference for Early Embryonic Morphogenesis of *C. elegans*
● Variability (Noise) of Division Timing, Division Orientation and Cell Arrangement
● A Conceptual Close-Packing Model for Cell Arrangement Up to 8-Cell Stage
● Quantitative Phenotyping Methods at Embryo and Cellular Level
● Cellular Phenotypes of 1818 Mutant Embryos (758 Genes) Before Gastrulation
● Categorized Phenotypes upon Gene Perturbation

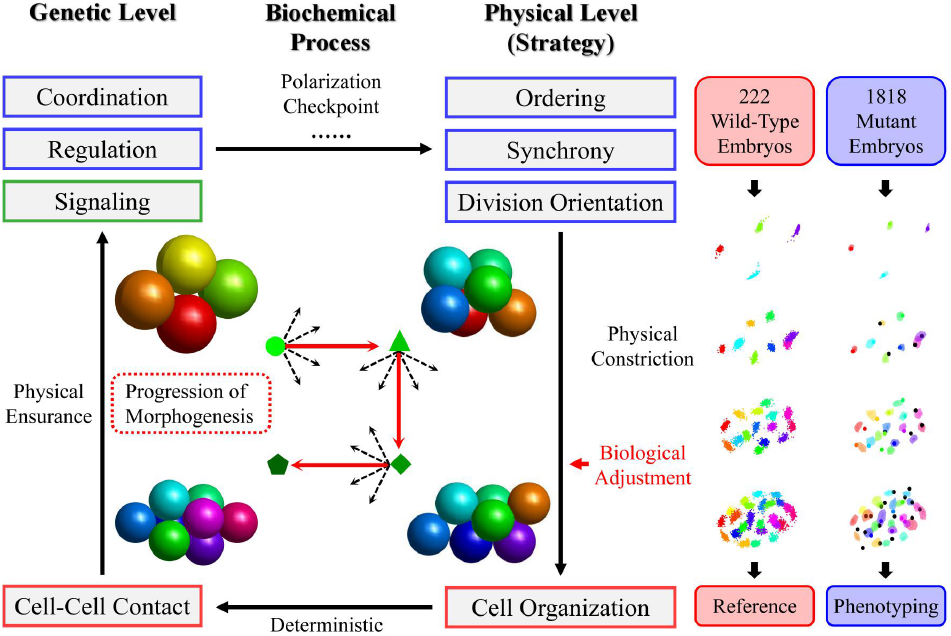

## Introduction

Morphogenesis is a spatio-temporal biological and physical process of multicellular structure formation, which is essential for signal transmittion, tissue specification and embryogenesis during development of metazoans, such as nematode^[1]^, fruit fly^[2]^ and zebrafish^[3]^. Failure in morphogenesis probably induces defective developmental procedure (e.g. cell-cell contact and interaction), leading to dysfunction, deformity, or even death in embryo^[4]^. Unusually, some small animals, like many species in nematode^[5]^ and gastrotrich^[6–7]^, have invariant cell lineage and constant cell number, which are classified as “eutely” creature. A recent study on genomes and morphology of four distinct nematode species proposed their conserved morphogenesis patterns maintained for over 20 million years^[8]^, which allow cells to have delicate intercellular signaling network and accurate differentiation at single-cell level^[1, 9–10]^.

*Caenorhabditis elegans*, a type of tiny transparent nematode which naturally lives in soil and has invariant cell lineage during embryogenesis, was widely used as model organism for developmental biology^[11]^, for that RNA-interference (RNAi) approach and automatic cell-tracing methods have been well developed and applied^[12–15]^, while its genome was also completely sequenced in Human Genome Project^[16]^. Most studies on early *C. elegans* morphogenesis paid high attention on the genetic and molecular mechanisms, for example, PAR proteins and Wnt signaling pathways have been investigated in depth and established as critical functional players for spindle formation as well as asymmetric segregation^[17–18]^. However, a lot of important details and problems still remain poorly understood as the following : how can cell position be organized so precisely and reproducibly among individuals? How robust or variable can the multicellular structure be? What are the regulatory roles of endogenous molecular activity and cell-level biophysical mechanics? To help answer these questions, several previous works provided wild-type databases for different developmental properties such as cell-cell contact, cell cycle, gene expression, differentiation and morphogenesis^[19–23]^. Statistical reference based on numerous wild-type embryos as well as quantitative mutant phenotypes of multiple genes are important keys to unravel the secrets of *C. elegans* embryonic morphogenesis.

In this paper, 222 wild-type *C. elegans* embryos were cultured and imaged using time-lapse 3-dimensional (4D) confocal microscope with time resolution of approximately 1.5 minutes. Automatic tracing, manual quality-control editing and quantitative analysis for cells before gastrulation onset (4- to 24-cell stage) generated spatio-temporal statistical variability of developmental properties for each cell. Further measurement on cell division timing and kinematic state revealed that 8 groups of cells could be clustered with invariant division order and cells could still be in motion when others start to divide, implying strong robustness against disturbance caused by cell movement. To uncover the internal regulatory mechanisms on morphogenesis of early *C. elegans* embryo, a total of 758 genes were selected for RNAi perturbation and phenotyping based on the wild-type reference, constructing a genetic architecture coordinating cell-arrangement progression over time at single-cell level. Causal relationships between defects of division timing, division orientation and cell arrangement were investigated and then an amount of mutant cases with different defective behaviors were found (e.g. anti-interference, induced defect and recovery from defect), uncovering redundant coordination as well as self-repairing ability in morphogenesis during *C. elegans* embryo development. Moreover, the wild-type reference was compared to a close-packing model for cell arrangement during 4- to 8-cell stage, which well accorded with the known essential cell-cell contact and signaling, suggesting high structural similarity and possible basic mechanical cues coordinating cell arrangement. Last but not least, a system-level phenotyping method and a public software were designed for looking over our wild-type reference, mutant phenotypes and automatically phenotyping new embryos inputted.

## Establishment of wild-type morphogenesis reference with spatio-temporal properties at single-cell level

To statistically construct a wild-type morphogenesis reference on cell variability with little bias or error, we first set up a pipeline mainly consisting of data acquisition, quality control, data processing and data integration (Fig.1A). Using experimental and computational methods described previously^[21]^, time-lapse 3D position of each cell from 4- to 350-cell stage was imaged, identified and traced in 222 wild-type *C. elegans* embryos (Fig.1B). All the wild-type imaging experiments were carried out at room temperature of 20-22℃, with temporal resolution of about 1.5 minutes and spatial resolution of 0.09 μm/pixel in focal plane and no more than 1 μm/pixel along shooting direction (Table S1). Before 350-cell stage, *C. elegans* cell lineage consists of 6 major sub-lineages : AB128, MS16, E8, C8, D4, Z2&Z3 (descendants of P4), while all the first 5 groups would continue to divide and Z2&Z3 remain undivided until postembryonic stage (Fig.1C, Fig.S1E)^[1]^. Thus, for each wild-type embryo, complete lifespans of AB4-AB128, MS1-MS16, E1-E8, C1-C8, D1-D4, P3 and P4 cells as well as division timing of AB2, EMS and P2 were recorded, in other words, all the cells of AB256, MS32, E16, C16 and D8 must appear before the ending time of imaging, producing a final state with approximately 350 cells in total, so as to establish a spatial-temporal reference for each cell with equal sample size. To eliminate editing error, all the cells with lifespan beyond their normal ranges (MEAN ± 3*STD, 222 wild-type samples) were taken out for manual examination and revision, to make sure that the 4D tracing was correct and the whole dataset possessed high credibility (Supplementary Material 1).

**Figure 1.**
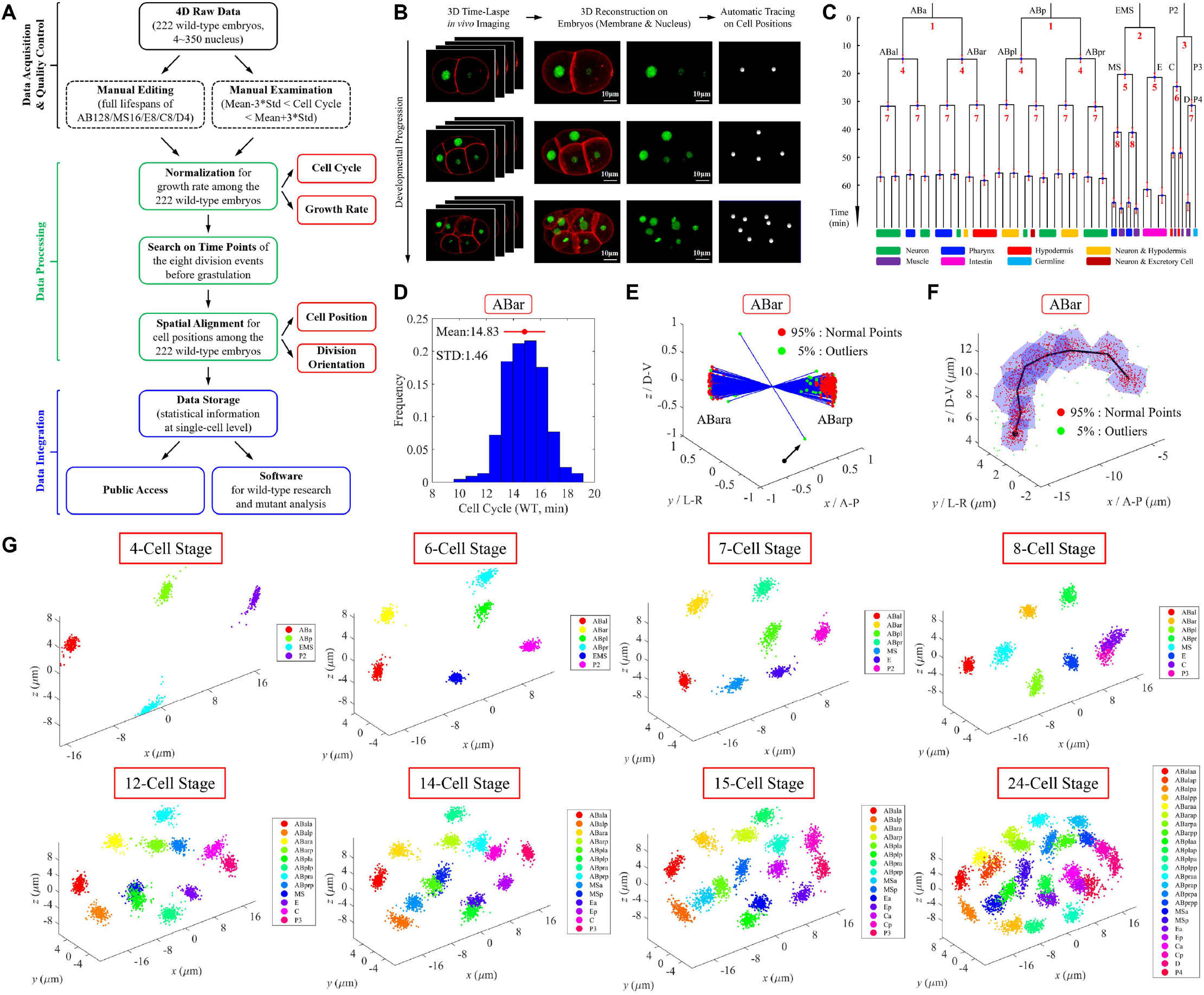
Establishment of wild-type morphogenesis reference with spatio-temporal properties at single-cell level. A. A pipeline consisting of data acquisition, quality control, data processing and data integration. B. Time-lapse 3D *in vivo* imaging, embryo reconstruction and automatic cell-position tracing on a *C. elegans* embryo expressing GFP in nucleus (green) and PH(PLC1d1) in membrane (red). The whole duration lasted from 4- to 350-cell stage; the membrane marker here is only for illustration purpose, as most of the data in this work was not obtained from this strain; 2-cell, 4-cell and 8-cell stages are presented (Supplementary Material 2). C. Cell-lineage tree up to 51-cell stage with tissue-differentiation information^[1,21]^ and cell grouping based on invariant division ordering in wild-type embryo (Table S4). Precusor cells of each lineage are marked with cell identity; Z2&Z3 are not shown here as they are generated after 51-cell stage and remain undivided until postembryonic stage; error bar denotes standard deviation (STD) of normalized cell cycle; developmental time is shown on left with a vertical axis, using the last moment of 4-cell stage as the origin. D. Cell-cycle distribution formed by 222 wild-type samples. Taking ABar as an example, histogram with average of 14.83 minutes and standard deviation 1.46 minutes is illustrated. E. Division-orientation distribution formed by 222 wild-type samples. Taking ABar as an example, initial distance between two daughters is normalized to one; 95% normal samples are plotted with red points while the other 5% outliers with green points; the extreme outlier indicated with black arrow was found to be an editing error. F. Migration-trajectory distribution formed by 222 wild-type samples. 95% normal samples are plotted with red points and a convex polyhedron with blue shade, while the other 5% outliers with green points; black line denotes the average migration trajectory, with a black point labeling the origin. G. Cell-arrangement distribution formed by 222 wild-type samples after spatial alignment from 4- to 24-cell stage (Fig.S2). Each color represents one specific cell identity, noted in legend on right; *x*, *y*, *z* axes correspond to anterior-posterior (A-P), left-right (L-R) and dorsal-ventral (D-V) axes respectively.

Using all the cells with complete lifespan, difference of growth rate between wily-type embryos were compared and normalized by proportional function, based on their good proportional relationship reported before (Fig.S1ABC, Table S1)^[24–25]^. The linear regression revealed a smallest *R*^2^ (goodness of fit) between any two embryos larger than 0.80. Besides, less than 1% of samples have relative global growth rate faster than 1.12 or slower than 0.85, which was regarded as the threshold of abnormal proliferation rate in our experimental conditions and used for mutant screening on division-timing defect (Fig.S1D).

In this work, we specifically chose 4- to 24-cell stage as developmental duration for morphogenesis study, for that maternal-zygotic transition and gastrulation haven’t been activated yet, so that the embryo system would be comparatively simplified^[4, 26]^. Right before the onset of gastrulation, cells could be classified into 15 groups based on known information about synchronous and asynchronous division timing as well as cell-fate differentiation in early *C. elegans* embryogenesis^[1, 27–30]^, which are AB2, AB4, AB8, AB16, EMS, E, E2, MS, MS2, MS4, P2, C, C2, P3, D and P4 cells (Fig.1C, Table S2). For example, cells differentiated from germline (P1 → P2/EMS, P2 → C/P3, P3 → D/P4, P4 → Z2/Z3) acquire asymmetric proliferation rate and cell fate, meanwhile Wnt signaling from P2 cell continues to induce EMS and its daughters (MS, E) differentiated^[18]^. On the basis of these grouping rules, cell positions at the first and last moment when a whole group exists before 24-cell stage were extracted from all the wild-type embryos, composing a temporally sequential matrix containing cell position information in 23 independent time points (**frames**). If average duration between two consecutive frames was longer than 3 minutes, it would be further segregated evenly, so that a temporal sequence with interval no longer than 3 minutes could be achieved, as a result, 6 more frames were inserted (Table S2). Subsequently, for each frame, global linear transformations including rotation, translation and scaling were performed on all the cell positions to minimize systematic variation between the samples due to eggshell size, compression by slide mounting, random cell movement inside the embryos, etc (Fig.S2).

Though reproducibility and variability of morphogenesis have been previously studied^[1, 23, 31]^, underlying mechanism still remains unclear due to lack of enough quantitative information from wild-type statistics and mutant phenotypic analysis. So far, with cell positions from 222 wild-type embryos normalized, a precise and reproducible 4D cell-arrangement pattern was reconstructed (Fig.1G). As well, three developmental properties at single-cell level were also statistically obtained : cell cycle, division orientation and migration trajectory. ABar cell, which has been proposed to shift posteriorly during establishment of left-right asymmetry and meet C cell to receive Wnt signal^[32]^, was taken as an example to show the cell-level reference (Fig.1DEF). Therefore, a quantitative and statistical framework of wild-type early morphogenesis has been established, laying the foundation for further systematic research on RNAi-perturbed mutant embryos as well as the normal ones.

As the cell positions of each cell group’s first co-existence moment were extracted, variability of division orientation was estimated by calculating the narrowest cone which encircles all the division vectors between two daughter cells, showing that orientations of all cell divisions before 24-cell stage were controlled within a half angle (i.e. maximum angular deviation from the rule vector, 99% samples) approximately smaller than 45° but vary from 16.05° (ABpra) to 45.45° (MS) (Fig.S4, Table S3), probably affected by PAR proteins, Wnt/Src-1 signaling and other biochemical regulations^[17–18, 33–36]^. On the one hand, different level of division-orientation accuracy potentially implied different biologically regulatory mechanisms as well as different mechanical requirement for cell division. On the other hand, well-tuned division orientation ensures morphogenesis directionally progress to its targeted structure according to expected developmental blueprint. By physical modeling and computational simulation, previous theoretical research suggested that change of division orientation may lead to misplacement of cells due to cell-cell and cell-eggshell mechanical interaction^[37]^. Under mechanically deterministic assumption, a naturally available variation limit on division orientation should exist and may be reached in RNAi-perturbed embryos, which could be evidence for further supporting or verifying this physical viewpoint.

A previous study also used similar experimental methods and acquired 53 wild-type embryos to establish *C. elegans* embryogenesis reference in multiple dimensions including morphogenesis^[23]^. Nevertheless, information of several cells before 8-cell stage was missing, probably due to difficulty on controlling the initial imaging time. Apart, only one time point of data was used for statistics on cell position, leading to lack of complete migration trajectory as well as division orientation. Another study on cell variability also built a wild-type *C. elegans* morphogenesis reference consisting of cell division timing (cell cycle), cell position and division orientation, but only collected 18 embryo samples^[25]^. In our reference, all the 222 wild-type embryos were cultured and imaged from 4-cell stage and reconstructed the early morphogenesis procedure at time resolution of less than 3 minutes, generating new statistical and complete cell properties (e.g. cell cycle, division orientation, migration trajectory) which exhibit lower variation than the previously published ones (Table 1AB, Fig.S3, Fig.S4).

**Table 1A.** Comparison between our wild-type reference and a previous one (Moore et al. 2013).

| Selected Properties of Morphogenesis Reference | Previous Reference<sup>[23]</sup><br>(Moore et al, 2013) | Our Reference | Brief Description of Our New Reference |
| --- | --- | --- | --- |
| Sample Number of Wild-Type Embryo | 53 | 222 | All the embryos contributed completely and equally. |
| Initial Valid Stage | 8-Cell Stage<br>(ABa/ABp/EMS/P2/MS cells are missing.) | 4-Cell Stage | All the embryos were imaged from 4-cell stage. |
| Division Orientation | / | √ | Division orientation of each cell was statistically formed by 222 vector samples. |
| Time Resolution | Only one time point was used.<br>("5 minutes into the cell's life") | Less than 3 Minutes | 29 frames of cell-arrangement structure were generated. |

**Table 1B.** Comparison between our wild-type reference and a previous one (Julia et al. 2013).

| Selected Properties of Morphogenesis Reference | Previous Reference<sup>[25]</sup><br>(Julia et al, 2013) |  | Our Reference |  |  | Brief Description of Our New Reference |
| --- | --- | --- | --- | --- | --- | --- |
| Sample Number of Wild-Type Embryo | 18 |  | 222 |  |  | All the embryos contributed completely and equally. |
| Division Orientation | Not published.<br>(ABa / ABp $\approx 60^\circ$)<br>(Mean Deviation of All the Samples) | | √<br>(ABa / ABp $\approx 30^\circ$)<br>(Maximum Deviation of the 99% Samples) | | | Tougher standard but smaller variability / noise. |
| Cell Cycle | Coefficient of Variation |  | Coefficient of Variation |  |  | The majority of cells have smaller variability / noise in cell cycle. The rest ones have similar values in both references |
|  | ABal | 0.329 | ABal | 0.094 | ↓ |  |
|  | ABar | 0.320 | ABar | 0.098 | ↓ |  |
|  | ABpl | 0.322 | ABpl | 0.094 | ↓ |  |
|  | ABpr | 0.327 | ABpr | 0.096 | ↓ |  |
|  | ABala | 0.121 | ABala | 0.081 | ↓ |  |
|  | ABalp | 0.122 | ABalp | 0.079 | ↓ |  |
|  | ABara | 0.115 | ABara | 0.083 | ↓ |  |
|  | ABarp | 0.103 | ABarp | 0.079 | ↓ |  |
|  | ABpla | 0.132 | ABpla | 0.091 | ↓ |  |
|  | ABplp | 0.114 | ABplp | 0.089 | ↓ |  |
|  | ABpra | 0.110 | ABpra | 0.084 | ↓ |  |
|  | ABprp | 0.115 | ABprp | 0.087 | ↓ |  |
|  | MS | 0.181 | MS | 0.086 | ↓ |  |
|  | MSa | 0.064 | MSa | 0.080 | ↑ |  |
|  | MSp | 0.063 | MSp | 0.081 | ↑ |  |
|  | E | 0.180 | E | 0.083 | ↓ |  |
|  | C | 0.089 | C | 0.088 | ↓ |  |
|  | P3 | 0.077 | P3 | 0.081 | ↑ |  |

## Robust progression of morphogenesis induced by division events with precise timing and coordinated ordering

Due to unequal segregation of cell-fate determinant^[36]^ and multiple cell-cell signaling such as Wnt signaling from P2 to EMS^[18]^ and the first *Notch* signaling from P2 to ABp^[38–39]^, cells in the early stage usually possess unique identity not only for for their spatial behavior but also for their distinguishable internal transcriptional profiling^[40]^. Consequently, for “eutely” *C. elegans* embryo, a cell must be placed in an accurate position, where its descendents subsequently undergo a series of programmed intercellular interaction and differentiation, then reach their targeted final locations with tissue-specific function at larva stage^[1, 41]^. In other words, mislocation of a cell or location swap between cells should not be approved of in the developmental process of *C. elegans* embryo.

As cell-position variability was statistically quantified (Fig.1FG), an intriguing questions could be brought out again and argued : what are the physical and genetic roles respectively on organizing a single, specific and precise cell-arrangement pattern? Many researches focused on genetic and molecular level and uncovered some essential biological mechanisms coordinating division timing, division orientation and cell arrangement (e.g. spindle checkpoint)^[42–44]^. In addition, few attention was paid on the cell-scale self-organization resulting from fundamental mechanical cues, which seemed to contribute to pattern formation with biologically required cell location and cell-cell neighbour relation as well^[29, 37, 45]^. Several experimental and theoretical studies suggested that, in cell scale, division timing and division orientation are crucial fail-safe mechanisms for correct cell arrangement. Enough duration between two consecutive ordered division events allows the system to relax and cells to reach their expected positions before the next divisions start, otherwise the system may fail to progress normally because it’s still in motion and unstable. Besides, precise division orientation could ensure the progression direction of the cell-arrangement structure in embryo^[29, 37]^.

However, those hypotheses lacked of direct and statistical verification of experiment. To elucidate the physical mechanism on robust morphogenesis of *C. elegans* embryo, cell behavior has to be comprehensively dissected on both dimensions of space and time. As well, one should evaluate 3 dimensions of system characteristics by experiment :

● How reproducible or variable is the wild-type multicellular structure?
● Are cell divisions always coordinated in distinct order with enough interval for relaxation?
● Do cells actually relax and reach their expected positions before the next divisions start? By inspecting the cell behaviors in experiment, the 3 questions will be answered one by one as below.

At first, we quantitatively evaluated the dynamics of cell-position variability. For each cell at a specific frame, 222 sample points’ distances (i.e. displacements) to their average positions 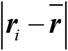 were used to calculate their standard deviation 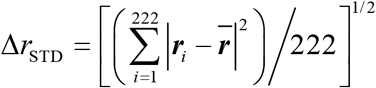 as a symbol of spatial variability (Fig.2AB). Globally, cell variability was slightly increasing over time (*R*^2^ = 0.390) but has deviation (STD) always smaller than 2.5 μm, while radius of cell in 24-cell stage is longer than 5.0 μm (i.e. Δ*r*_STD_ < *r*Cell)^[46]^. However, cells may have very different variability from each other, for example, at the moment of EMS division E cell had a much larger variable area than the other cells, probably due to its unstable and active movement, as EMS and E cells keep receiving Wnt signaling from P2 cell which coordinates the active rotation of MS-E cell pair before cytoplasmic separation^[18, 47]^. Whereas standard deviation Δ*r*_STD_ of cell position could represent the statistical area of how distant a cell could be shifted, it’s also important to consider the most extreme cases, so as to check the natural limit of system noise and robustness. To this end, box plot was carried out to highlight the mild outlier as well as extreme outliers (Fig.2C). For each cell, positions in over 75% wile-type embryos deviated within a radius of 2.5 μm, but there’re some extreme outliers shifting even farther than 8 μm which didn’t cause embryo death in the end, indicating strong tolerance on cell-arrangement noise in developing *C. elegans* embryo. Dynamic change of spatial variability was also connected to the kinematic state of system, which is lower during division progressing and getting higher later, exhibiting fluctuation in accordance with start-up times of division events (Fig.S5). In other words, cell division progresses in a relatively stable environment and after that spatial variability increases until the next divisions taking place.

**Figure 2.**
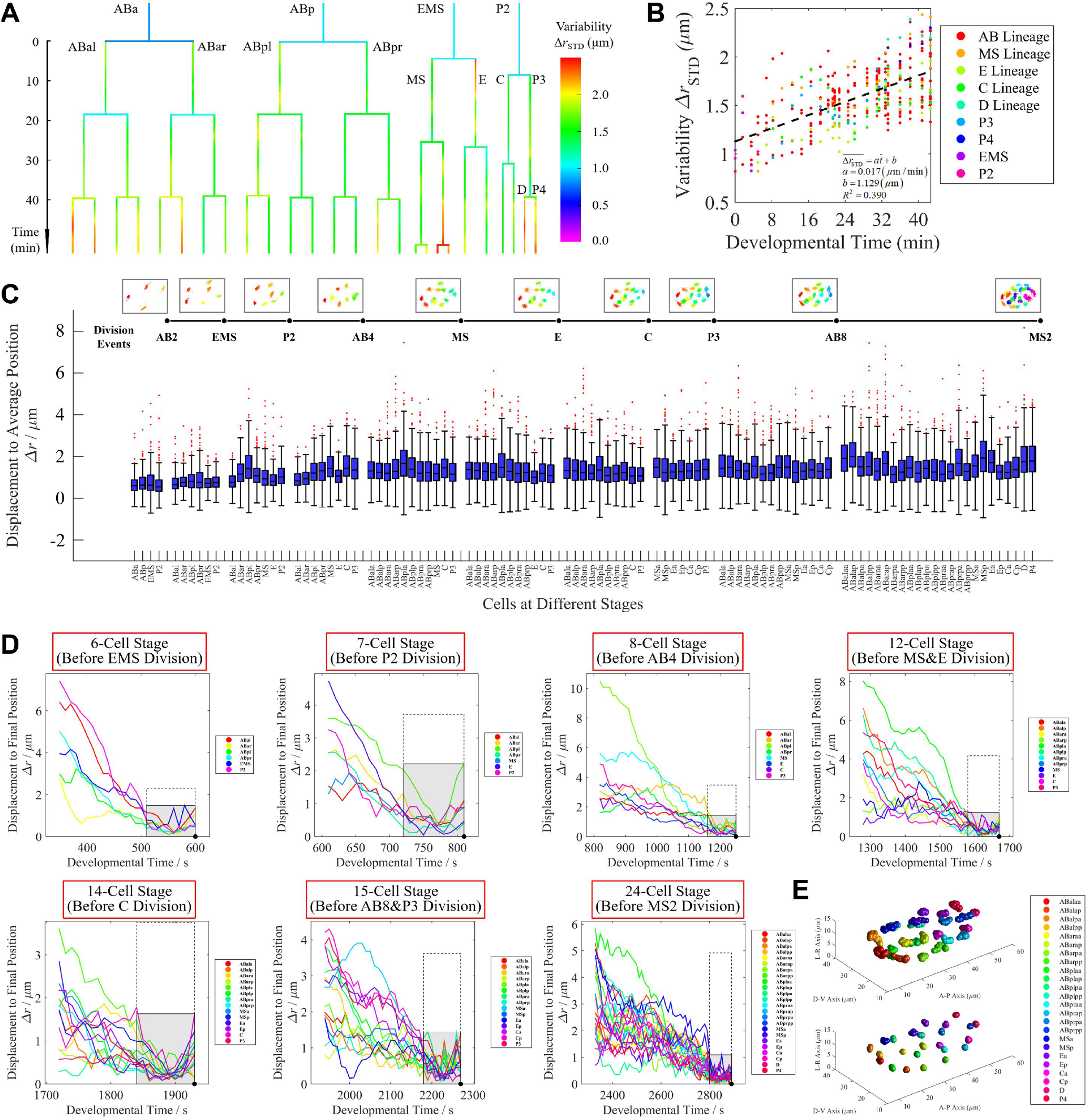
Spatial variability and movement of cells from 4- to 24-cell stage. A. Cell-lineage tree with spatial variability Δ*r*_STD_. Standard deviation Δ*r*_STD_, which is calculated with 222 sample points for each cell, is quantitatively illustrated with different color according to the color bar on right; developmental time is shown on left with a vertical axis, using the last moment of 4-cell stage as the origin. B. Increasing trend of spatial variability Δ*r*_STD_ of cells from 4- to 24-cell stage. Using data of all the 29 frames, spatial variability of different cell at each moment is plotted with different colors, according to corresponding relationships in legend (Table S2); linear regression reveals slope *a* of 0.017 μm/min, intercept *b* of 1.129 μm and determination coefficient *R*^2^ of 0.390. C. Distribution of positional deviation Δ*r* (i.e. displacement to average position) before division starting, formed by the 222 wild-type embryo samples. Moments before the 10 division events (AB2, EMS, P2, AB4, MS, E, C, P3, AB8 and MS2) are shown successively from left to right according to their average division time (Table S2); identities of cells existing in each moment are marked on horizontal axis, and distribution of their positional deviations Δ*r* is illustrated using box plot. D. Distance of cell to its final position (i.e. displacement to average position in the last 1.5 minutes before division) Δ*r*. Black point represents initial timing of the next division; gray area surrounded by solid lines represents the maximum displacement in the last 1.5 minutes, while the white area surrounded by dashed lines represents the maximum normal deviation Δ*r*_Q3+1.5*IQR_ (Upper Quartile + 1.5*Interquartile Range). E. Cell movement trajectory in the complete duration (upper) and the last 1.5 minute (lower) of the 24-cell stage. Each color represents one specific cell identity, noted in legend on right.

For the second question, if cell really needs enough time for relaxation *in vivo*^[29]^, their division order and relaxation duration should be reexamined experimentally. Despite of any known information on symmetry breaking by division^[1, 26–30]^, here we used the division-timing data from 222 wild-type embryos to search the ordering rules with generality. Statistical analysis on division timing of cells from 4- to 24-cell stage found a set of elaborate temporal-sequential coordination, that is, all the cells are either in distinct order or synchronously coupled together (Fig.1C). Here, 4 sets of multicellular division events (average maximum interval shorter than 1.5 minutes and no invariant order existing between any two cells) and 6 pairs of division events kept in invariant order (average interval longer than 3 minutes) with probability of 100% were concluded from 222 wild-type embryos (Fig.S6, Table S4). Note that E and MS were clustered together because MS usually divides only about 1 minute earlier than E due to differentiation (Table S2B)^[1, 26, 48]^. As well, AB8 and P3 were found to divide together, possibly responsing to the same signal pathway undiscovered. For each division event consisting of more than one cells, no invariant order was found between any two of them, suggesting that division timing of cells within a group was commonly regulated to be synchronous with slightly random noise. Even though cells were clustered into 8 groups dividing in distinct order, duration between two successive groups varies widely among wild-type embryos. For instance, interval between AB2 and EMS division ranges from 1.23 to 8.83 minutes (Table S2B). This high variation implied strong system robustness against temporal and motional noise again, for that cells may not have enough time to reach a stable state before onset of the next division event. It’s worth noting that, numerous wild-type samples enable us to find the extreme natural cases and establish common conclusions more statistically and precisely.

So far, both spatial and temporal variability revealed strong robustness against dramatic intrinsic motional noise. The fail-safe mechanical model for self-organization proposed before^[29]^ may not be able to explain this outstanding feature. To comprehensively figure out if these outstanding coordination rules on cell division timing (i.e. ordering and simultaneity) really contribute to robustness and precision during embryo morphogenesis, one needs to look deep into cell behavior at the interval between two consecutive division events. For this purpose, imaging with time resolution of 10 seconds was performed on a *C. elegans* embryo from 4- to 24-cell stage, until the division of MS2 cells, to help inspect whether the cells use an interval long enough for relaxation before the next cell division events (Supplementary Material 3). In the last 1.5 minutes of each stage, cells have reached and stayed in their wild-type normal regions (Δ*r* < Δ*r*_Q3+1.5*IQR_) (Fig.2CD). Compared with cell movement in the complete duration between two division events, the cells kept relatively still at their final positions in the last 1.5 minutes just before the next division event, with little motional variability (Fig.2DE). For majority of stages (6-, 12-, 24-cell stages, all cells), a cell has a much smaller motion velocity in the last 1.5 minutes than the one in the first 1.5 minute within a stage, which means the embryonic system moves dramatically as division starts up and then relaxed gradually with slow movement eventually (*v* < 2 μm/min)(i.e. velocity changed from high to low) (Fig.S7). As all of these stages are driven by division of AB cells, which occupy 1/2 ∼ 2/3 of cells in embryo and always divide synchronously, this mechanical and global perturbation could be a reasonable explanation for the relaxation phenomenon, supporting the clock-setting viewpoint proposed before^[29]^. However, at the other stages (7-, 8-, 14-, 15-cell stages), a cell may also acquire higher velocity compared to its initial state (*v* > 3 μm/min), indicating that divisions of the other cells with small number (e.g. EMS, P2, MS, E, C) may progress in a unstable and noisy embryo environment, likely due to molecular active motion as well as lack of time for relaxation (Fig.S7). In a word, duration between division events indeed probably provides time for cell motion and system relaxation, so that the next division events could start up at a relatively steadier environment. Despite all this, sometimes cells may still move speedily when a group starts to divide, for example, in the 7-cell stage ABpl moved directionally away from its average position in the last 1.5 minutes (*v*_ABpl_ ≈ 3.35 μm/min), which also has the longest travelled distance as well as the largest spatial variability compared with other AB cells (Fig.2D, Fig.S5, Fig.S7, Table S5)^[25, 32]^. Migration accuracy of cell in those motional stages could probably achieved by active cell motion and reorientation, which needs to be further investigated with biological experiments (e.g. mutant screening)^[47]^. Owing to non-ignorable photoxicity and lethality, time resolution and imaging duration of this experiment are limited, so that subtle movement (within interval of 10 seconds) in details may be missing.

## A genetic architecture coordinating early C. elegans morphogenesis spatially and temporally

On the basis of the established wild-type reference on cell cycle, division orientation and migration trajectory before onset of gastrulation, namely 24-cell stage, system-level and cell-level defect screening was performed on 1818 RNAi-perturbed mutant embryos according to a pipeline designed (Fig.3A). A total of 758 genes were selected based on their degree of conservation and reported phenotypes upon perturbation, and at least two replicates were curated for each gene (single-gene RNAi perturbation) (Table S6)^[21]^. All the mutant embryos were manually curated up to approximately 350-cell stage, except for the ones with too dim expression marker signal or unusually slow proliferation pace, cell-cycle arrest and apoptosis phenotypes (i.e. the 350-cell stage in those mutants may be beyond the ending time point of imaging)^[49]^. Similar to the wild-type embryos, cell positions of mutant ones at time points before or after divisions were extracted according to the time-setting methods mentioned above (Table S2), then normalized and compared to the wild-type reference (Fig.S2). For test on division-timing defect, cell cycles of cells in a mutant embryo would be used for proportionally linear fitting to their wild-type averages group by group successively based on their division order (Table S2, Table S4), if any cell was unable to be normalized to its confident interval formed by 98% of wild-type samples (1% for each side to represent the abnormally fast and slow division rate) no matter what value was adopted to the global scaling parameter, this cell group would be taken as the first cell and timing that division-timing defect occurs (Fig.3B). The fitted scaling parameter acquired, namely relative growth rate, was then compared to the wild-type distribution to estimate the mutant effect on global proliferation pace (Fig.S1D). For each cell’s position, 99% of wild-type sample points nearest to their average were selected out to form a convex surface, then the cell located outside the convex polyhedron would be regarded as a defective outlier, so as to estimate defect of cell division orientation and cell arrangement at single-cell level. For all the three cell-level properties, namely division timing, division orientation and cell arrangement, if a cell was detected with abnormality not only at the same time point but also in at least two embryo replicates, it would be taken into account as a reproducible defect. The severity of embryonic cell-arrangement defect was quantified by percentage of misarranged cells at each stage (Fig.3E). The screening procedure ensured false positive of each single-cell property smaller than 1%, while over 80% of wild-type embryos could completely passed the test without any screened abnormality (Fig.S8)(Supplementary Material 4).

**Figure 3.**
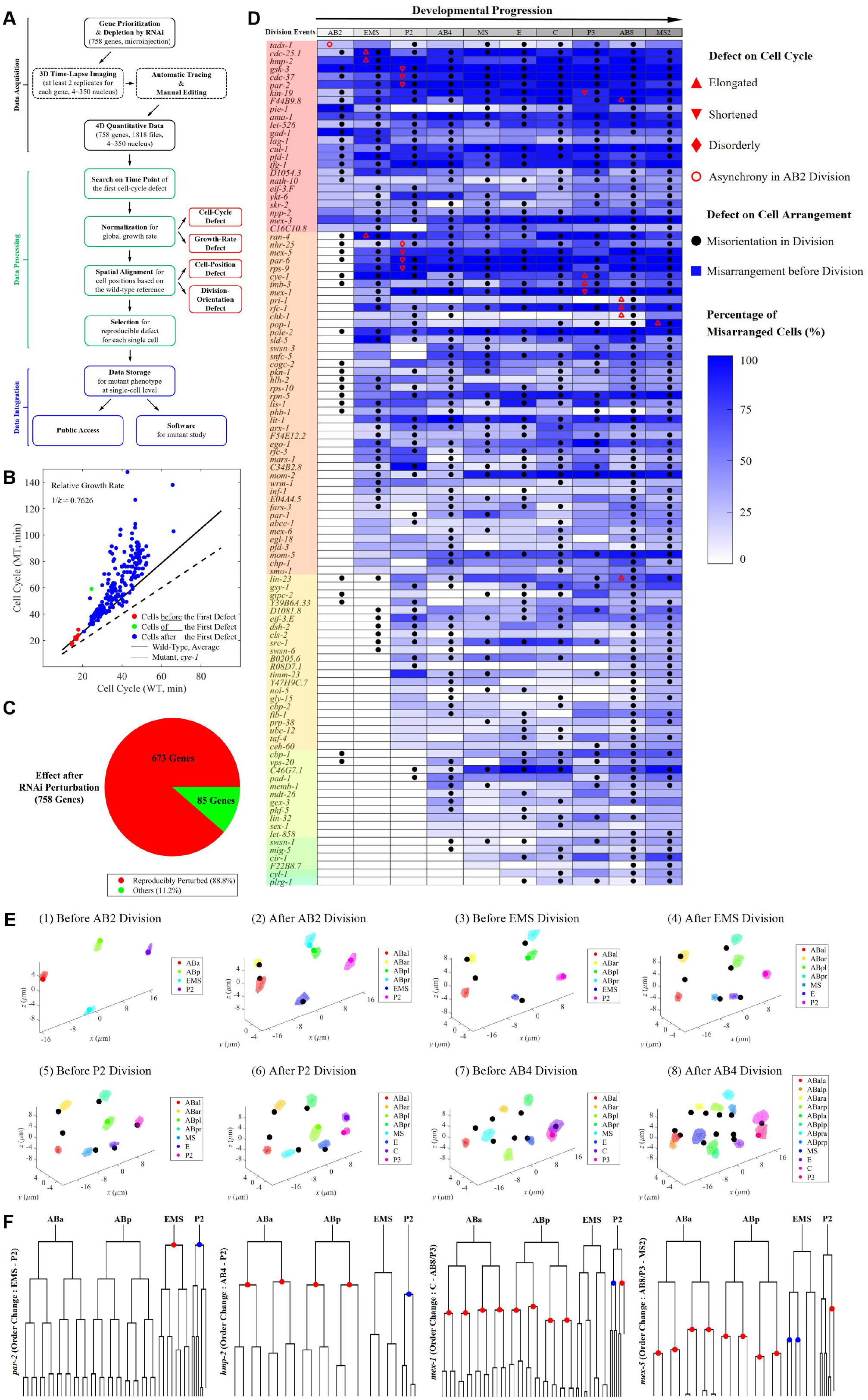
Defect detection on RNAi-perturbed mutant embryos. A. A pipeline consisting of data acquisition, data processing and data integration. This phenotypic analysis provided several dimensions of morphogenesis defect, including global growth rate, cell cycle, division orientation and cell arrangement. B. Normalization on global growth rate, detecting and providing the linear region as well as the first cell-cycle defect. C. Sector diagram on reproducible perturbation in the mutant embryos. Mutants of 673 genes showed reproducible perturbed phenotype, while the others are unreproducible or completely not effected. D. Reproducible results of defect detection on RNAi-perturbed mutant embryos. Only those with reproducible phenotype and at least 25 percent of cells misarranged at 24-cell stage are shown; the 10 division events (8 stages) are listed successively along the lateral axis on top; the genes are listed along the vertical axis on left based on their initial timing of cell-arrangement defect (noted in panel with different colors); blue shaded rectangle with different darkness represents the percentage of misarranged cell; red triangle, diamond and circle represent different types of cell-cycle defects; black point represents defective division orientation; “disorderly” means cell cycle of a cell was significantly elongated as well as shortened in different RNAi samples of the same gene. E. Defect detection on cell position as well as division orientation, using a *mex-5* RNAi-perturbed embryo (serial number 217, Table S6) as example here. Each color represents one specific cell identity, noted in legend on right; misarranged cells in mutant are illustrated with black point while the normal ones with original color according to the legend. F. Change of division order in mutant. From left to right : *par-2*, EMS → P2 synchrony restoration; *hmp-2*, AB4 → P2 order reverse; *mex-1*, C → P3 synchrony restoration; *mex-5*, AB8/P3 → MS2 asynchrony breaking.

In our examination scope, which is 4- to 24-cell stage, 88.8 % of genes perturbed showed reproducible defects, while the other 11.2% are unreproducible or not perturbed at all, probably due to insufficient samples, unstable and unreproducible perturbed behaviors, potential experimental failures, or naturally inactivated state before zygotic gene activation (Fig.3C, Table S6). With defect information on division timing, division orientation and cell arrangement, a total of 106 genes with at least 25% of cells misarranged at 24-cell stage were selected out to present a genetic architecture coordinating early *C. elegans* morphogenesis successively (Fig.3D).

The screening results were similar to our expectation, for example, maternally provided PAR proteins (*par-1*, *par-2*, *par-6*), which were found to regulate timing of mitotic entry, rate of DNA replication, cell polarity and asymmetric segregation, RNAi perturbation of those proteins could lead to serious early disorder in many aspects (Fig.3D)^[17, 34–35, 50–51]^. Genes with severe mutant defect also included other functional factors involved with Wnt signaling (e.g. *mom-2*, *mom-5*, *kin-19*)^[18, 33]^, Notch signaling (e.g. *lag-1*)^[38, 39]^, E3 ligase (e.g. *cul-1*, *skr-2*, *lin-23*)^[21]^, RNA splicing (e.g. *D1081.8*, *prp-38*, *let-858*)^[26]^, etc (Fig.3D). Interestingly, five types of reproducible changes in division order were found in genes as follow : EMS → P2 (*par-2*, *par-6*), P2 → AB4 (*hmp-2*), MS & E → C (*par-2*, *par-6*), C → AB8 & P3 (*mex-1*, *hmp-2*, *ran-4*, *kin-19*, *cdc-25.1*), AB8 & P3 → MS2 (*mex-1*, *mex-5*, *cye-1*, *gsy-1*), which will all lead to serious misarrangement of cells and death in embryo (Fig.3DF, Fig.S9). These screening results not only systematically quantified the mutant phenotypes, but also revealed the initial timing on morphogenesis defect, the identity of defective cell and the potential function of genes, thus, providing new layers of information for early morphogenesis and embryogenesis..

Overall, cell arrangement usually goes wrong along with defect of division orientation, because both properties are spatially based on nucleus position from imaging experiments and totally same in essence. Concerning the causality between division orientation and cell arrangement, some cases such as *ran-4*, *nhr-25* and *mex-5* mutant during 4- to 6-cell stage, start from a normal cell-arrangement state with defective division orientation and quickly reach a defective cell arrangement (Fig.3DE). On the contrary, another cases such as *rps-9*, *mex-1* and *pri-1* mutant during 4- to 6-cell stage, start from a normal cell-arrangement state with correct division orientation of AB2 cells, but result in defective cell-arrangement pattern before EMS division (Fig.3D, Fig.S10). This ambiguous phenomenon is also contradiction between roles of Newtonian mechanics and active molecular regulation on embryonic morphogenesis. Briefly, some perspectives from physicists suggest that simple cell-level mechanical rules (e.g. clock setting, accurate division orientation and intercellular elastic pressure) ensure the cell organization before gastrulation^[29, 37, 45]^, while some biological experiments revealed that active cell motion is critical for correct pattern at early stage, such as reorientation of cell division axis due to cortical non-muscle myosin flow^[47, 52]^. To thoroughly elucidate the regulatory mechanism on morphogenesis and provide valid experimental results for different hypotheses, different kinds of abnormal developmental behaviors must be generated with genetic perturbation and analysed statistically.

## Systematic phenotypic coding method for analysis on early morphogenesis

To systematically interpret the early morphogenesis phenotype of mutants, the three dimensions of state property were continually adopted to evaluate the embryonic defect : division timing *T_i_*, division orientation *O_i_* and cell arrangement *A_i_*, where *i* indicates the division event based on statistical ordering of the 10 events before gastrulation, which successively are AB2(1), EMS(2), P2(3), AB4(4), MS(5), E(6), C(7), P3(8), AB8(9) and MS2(10) (Fig.4A, Table S2). Note that even though MS,E and P3,AB8 respectively have very synchronous division timing in wild-type samples (Fig.S6, Table S4), they are also regarded as different division events in this coding method for that these cells were supposed to have entirely different cell fate and usually perturbed in mutant (Fig.3D, Fig.S9). Besides, MS is known to divide slightly faster than E^[1, 26, 48]^. As a result, the early embryonic morphogenesis before 24-cell stage was segmented into 10 stages based on division events and their statistical start-up time.

**Figure 4.**
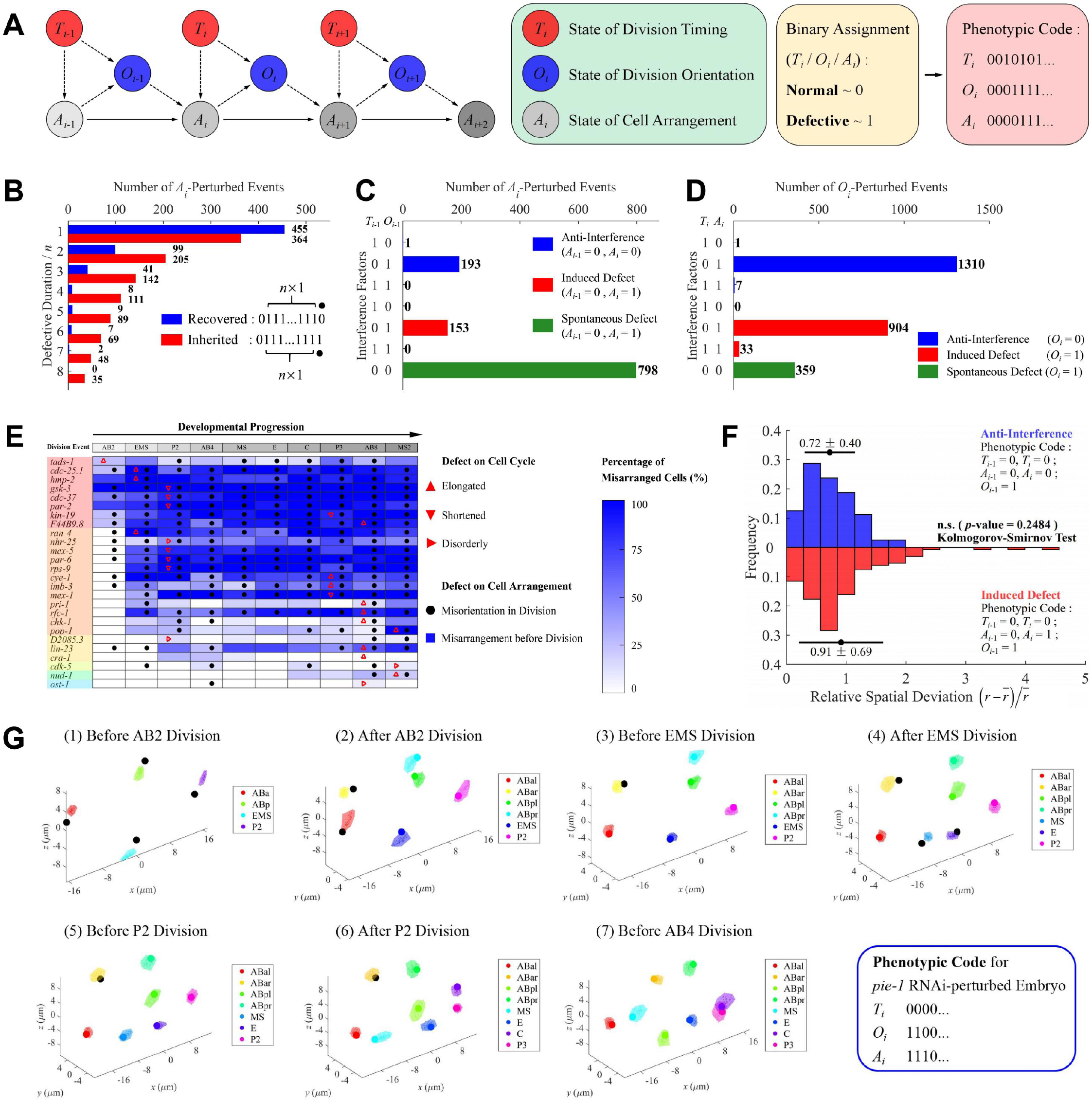
Phenotypic coding on mutant embryos reveals diverse morphogenesis behaviors. A. A systematic phenotypic coding method based on three state properties. Division timing, division orientation and cell arrangement are used to describe the state of early morphogenesis; each property was assigned 1 if defective, otherwise 0. B. Statistics on reproducibly recovered and inherited case after perturbation. Blue column denotes recovered ones with *A_i_* code 0111…1110; red column denotes inherited ones with *A_i_* code 0111…1111; number of cases is labeled next to the column; continuous defective stages (i.e. number of division events) *n* is labeled on the vertical axis. C. Statistics on different reproducible behaviors of cell arrangement responsing to division-timing and division-orientation perturbation. Blue column denotes anti-interference ones (*A*_*i-*1_ = 0, *A_i_* = 0); red columns denotes induced defect ones (*A*_*i-*1_ = 0, *A_i_* = 1); green columns denotes spontaneous defect ones (*T*_*i-*1_ = 0, *O*_*i-*1_ = 0, *A*_*i-*1_ = 0, *A_i_* = 1); number of cases is labeled next to the column; different combinations of interference factors (*T_i_*, *O_i_*) are labeled on the vertical axis. D. Statistics on different reproducible behaviors of division orientation responsing to division-timing and cell-arrangement perturbation. Blue column denotes anti-interference ones (*O_i_* = 0); red column denotes induced defect ones (*O_i_* = 1); green columns denotes spontaneous defect ones (*T_i_* = 0, *A_i_* = 0, *O_i_* = 1); different combinations of interference factors (*T_i_*, *A_i_*) are labeled on the vertical axis; number of cases is labeled next to the column. E. Reproducible results of defect detection on RNAi-perturbed mutant embryos, only those with reproducible division-timing defect are shown. The 10 division events (stages) are listed successively along the lateral axis on top; the genes are listed along the vertical axis on left based on their initial timing of cell-arrangement defect (noted in panel with different colors); blue shaded rectangle represents the percentage of misarranged cell, while red triangle represents defective cell cycle and black point represents defective division orientation; “disorderly” means cell cycle of a cell was significantly elongated as well as shortened in different RNAi samples of the same gene; “elongated” in AB2 cells (*tads-1*) means abnormal asynchrony in their divisions. F. Comparison between two groups of defective division orientation (anti-interference and induced defect) using relative spatial deviation 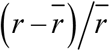 No significant difference was observed (Kolmogorov-Smirnov test, *p* = 0.3879). G. Cell-arrangement and division-orientation progression of *pie-1* RNAi-perturbed embryo (serial number 8, Table S6) during 4- to 8-cell stage, revealing recovery behavior from morphogenesis defect. Each color represents one specific cell identity, noted in legend on right; misarranged cells in mutant are illustrated with black point while the normal ones with original color according to the legend.

For each stage examined, the three state properties would be assigned 1 if it’s defective, otherwise 0, which provides a phenoypic code for each mutant at single-cell level (Supplementary Material 5). Taking AB4(4) division event as an example, if cell cycle of any one cell among AB4 were outside the confident range, *T*_4_ would be assigned 1; also, if any daughter of AB4 was spatially located outside its confident area at the first co-existence moment of AB8 cells, *O*_4_ would be assigned 1; as well, if any cell was spatially located outside its confident area at the last co-existence moment of AB4 cells, *A*_4_ would be assigned 1 (Supplementary Material 5).

In many biological cases, self-organized cells or tissue could recover themself from injury or perturbation^[53–54]^. Here we investigated several remarkable phenomenons including self-repairing ability and interference causing cell-arrangement defect. Firstly, only considering the cell-arrangement state property *A* for simplicity, if an embryo starts to be defective from the *i*^th^ event (*A_i_* = 0) and maintains the deformity compared to wild-type reference, but be able to totally recover after *n* division events (*A*_*i*+*n*+1_ = 0), it’ll possess a phenotypic code of *A* as 011…110 from *i* to *i* + *n* +1. On the contrary, if the defect is inherited, i.e., the embryo keeps significantly abnormal state for *n* + 1 events (*A*_*i*+1_, *A*_*i*+2_,…, *A*_*i*+*n*+1_ = 1), its phenotypic code of *A* should be 011…111 from *i* to *i* + *n* +1. Thus, all the reproducibly recovered and inherited cases in RNAi-perturbed mutants of 758 genes were screened and counted (Fig.4B). 455 cases were found to recover after experiencing one division event while 364 cases kept defective (*n* = 1). As division event number or defective duration increases, less and less recovery or inherited cases were found, but always there were more inherited cases than recovery ones in our database except the condition that the cell-arrangement defect only last for one division event (*n* = 1). These results supported natural self-repairing ability against genetic perturbation during *C. elegans* early morphogenesis. However, the self-repairing ability is also limited if some key genes are perturbed or the embryonic defect is too severe, for example, *pie-1* mutant (a maternal CCCH-type zinc finger protein functioning in germline-fate specification^[55]^) could completely recover at 8-cell stage but soon goes wrong as development progressing (Fig.3D, Fig.4G).

Mechanical cues such as division orientation have been proposed to drive targeted morphogenesis of cells or tissue^[56–57]^. From physical and mechanical perspective, the 3 state properties probably exhibit internal cause-effect relationship (hypothesis) that is, well-tuned division timing *T_i_* ensures the system could have enough duration relax to its correct cell-arrangement pattern *A_i_* before the next division *O*_*i*+1_ starting^[29, 37]^. As spindle formation and cell segregation need time for molecular reaction and also depend on the cell-arrangement pattern where they occur, state of division orientation *O*_*i*+1_ maybe relies on the other two properties, which also directly affect the next cell-arrangement pattern (Fig.4A). It’s worth noting that the data merging method and all the analysis are independent on any cause-effect hypothesis between the 3 state properties, so that similar analysis could be applied to any other biologically possible correlation as well.

Next, we sought to investigate whether division timing and division orientation are direct interference factors that cause cell-arrangement defect or not. *A_i_*-perturbed events in different (*T*, *O*, *A*) codes were classified and counted. Here, three remarkable behaviors of cell-arrangement state were defined, which are anti-interference (*T*_*i*-1_ + *O*_*i*-1_ ≠ 0, *A*_*i-*1_ = 0, *A_i_* = 0), induced defect (*T*_*i*-1_ + *O*_*i*-1_ ≠ 0, *A*_*i-*1_ = 0, *A_i_* = 1), spontaneous defect (*T*_*i*-1_ = 0, *O*_*i*-1_ = 0, *A*_*i-*1_ = 0, *A_i_* = 1), respectively correlated to state property combination of *T*_*i*-1_ and *O*_*i*-1_ as perturbation inducer (Fig.4C). Interestingly, in most of the cases (798), the defects are spontaneous, which means a number of cell-arrangement defects occur with neither previous division-timing nor division-orientation defect. Besides, nearly no cell-arrangement defect was found to be induced by abnormal division timing (*T*_*i*-1_ = 1, *A*_*i-*1_ = 0, *A_i_* = 1). Most division-timing defects occur along with or after cell-arrangement defect (e.g. *gsk-3*), while some others have no impact on the later cell arrangement (e.g. *cra-1*) (Fig.4E). As well, sometimes division-timing defect occurs with slightly abnormal cell arrangement simultaneouly, which could quickly recover before the next division (e.g. *D2085.3*, *ost-1*).

On the other hand, 193 division-orientation defects are followed by defective cell arrangement (*T*_*i*-1_ = 0, *O*_*i*-1_ = 1, *A*_*i-*1_ = 0, *A_i_* = 1) and 153 followed by completely normal one (*T*_*i*-1_ = 0, *O*_*i*-1_ = 1, *A*_*i-*1_ = 0, *A_i_* = 0)(Fig.4C). Using all the cases of anti-interference (*T*_*i*-1_ = 0, *T_i_* = 0, *O*_*i*-1_ = 1, *A*_*i-*1_ = 0, *A_i_* = 0) and induce defect (*T*_*i*-1_ = 0, *T_i_* = 0, *O*_*i*-1_ = 1, *A*_*i-*1_ = 0, *A_i_* = 1) among the 1818 mutant embryos, severity of defective cell division orientation was quantified with relative spatial deviation 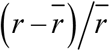. Comparison analysis showed no significant difference between two groups (Kolmogorov-Smirnov test, *p* = 0.3879)(Fig.4F). In other words, no inevitable relationship between division-orientation defect and the following cell arrangement according to our mutant-defect database. It’s worth pointing out that the division orientation here is defined as positions of daughter nucleus at their first moment, active reorientation or passive motion later is not involved here, which would probably play a key role in cell-arrangement correction as fail-safe mechanism^[47]^.

Similar statistical analysis was also performed on division-orientation defect, namely *O_i_*-perturbed events, using state property combination of division timing and cell arrangement as potential interference factors (Fig.4D). Expectedly, a total of 937 cases were found to have division-orientation defect along with cell-arrangement defect (*A_i_* = 1, *O_i_* = 1) because both of them are spatial variables and cell division orientation may inherit the defective cell-arrangement state and fail to maintain its normal orientation. Nevertheless, 1310 cases with abnormal cell arrangement followed by completely correct division orientation may indicate underlying biological mechanism that can rescue the perturbed process of spindle formation. Last but not least, 359 cases are those cell division orientation spontaneously goes wrong while the cell divides at normal time and position, implying these genes’ potential function in spindle formation and cell division.

Even though the analysis results depend on the mutant genes selected and didn’t give out much positive or detailed statistical conclusion here, they actually provided many different remarkable phenomenons, which are highly valuable for many developmental research topics, for example, the spontaneous cell-arrangement defect maybe results from perturbed active cortical myosin flow^[47, 52]^.

## A conceptual close-packing model reconstructs the cell-arrangement pattern during 4- to 8-cell stage

Cell position before gastrulation has been proved to be highly precise and reproducible among individuals by quantitative experiment of 222 wild-type embryos (Fig.1G, Fig2ABC). Configuration of cell arrangement ensures specific cell-cell contact which is required for cell-fate specification^[18, 38–39]^, division axis orientation^[47, 57–58]^ and other developmental procedure, in terms of both biology and physics. An overdamped mechanical model has been proposed to reconstruct the cell movement during 4- to 12-cell stage, using linear repulsive force to describe cell-cell and cell-eggshell interaction, which could well fit the results from both simulation and experiment^[37]^. To investigate the fundamental topology of cell-arrangement pattern, we further simplified the model by eliminating the difference of cell volume and elastic deformation, i.e., regarding a cell as incompressible sphere with even radius. Referring to the statistical wild-type reference, we manually reconstructed the multicellular structure based on close packing as well as major-and-minor axis asymmetry of ellipsoidal embryo, and then carried out normalization and comparison between them (Fig.5, Row 1-5). Cells from conceptual model and experiment with the same identity are always close to each other, with centroid distance *d* shorter than the radius of sphere *r*_s_, except for ABpr at 7-cell stage (*d* / *r*_s_ ≈ 1.05), because all the AB4, MS and E cells are the third generation of cells with similar volume, while P2 is still a second-generation cell, leading to a much larger volume for P2 cell than the others and giving rise to postitional shift of ABpr cell^[28,30,46]^.

**Figure 5.**
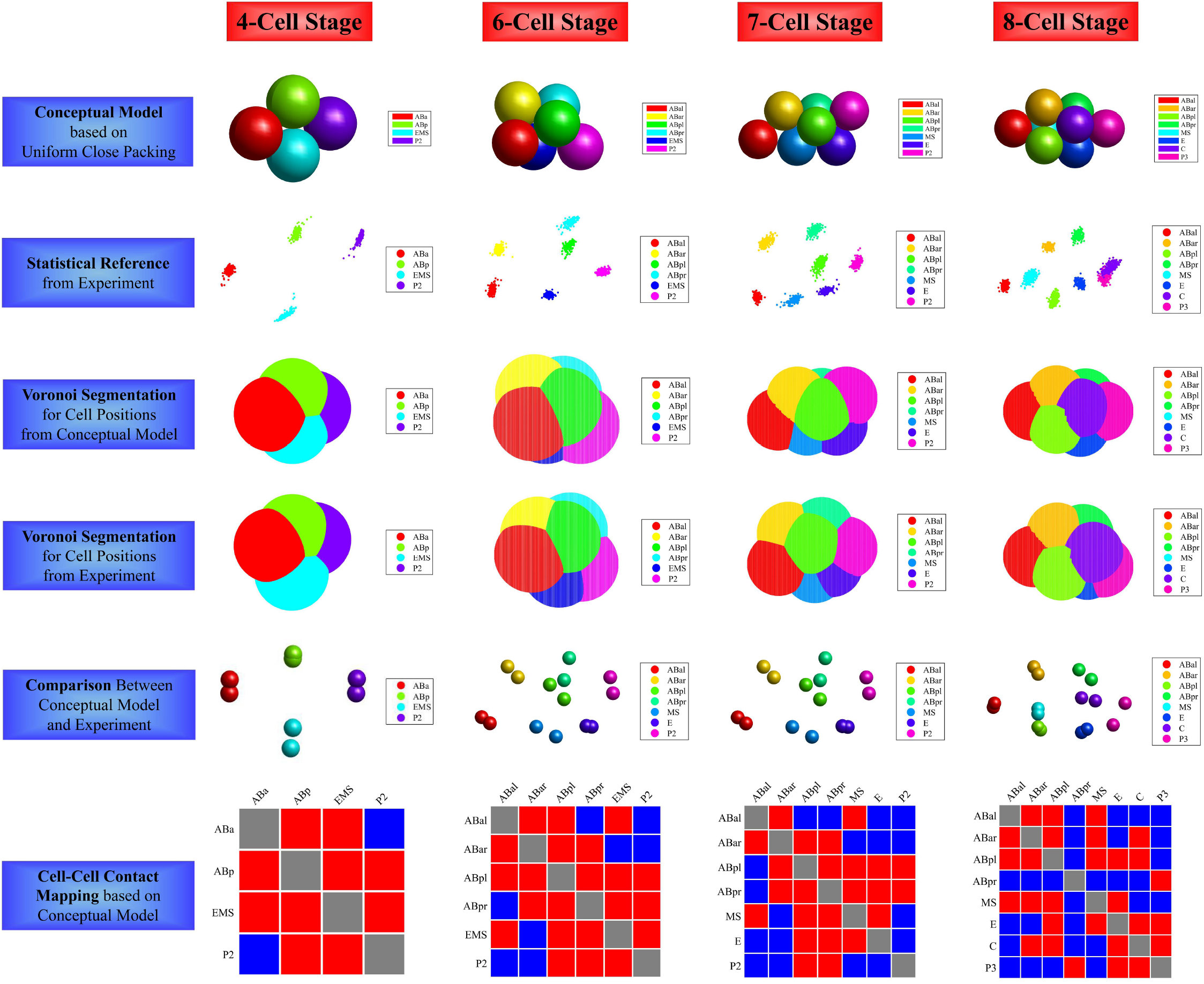
A conceptual close-packing model compared with cell-arrangement pattern from wild-type embryos. Only 4-, 6-, 7-, 8-cell stage are illustrated; each color represents one specific cell identity, noted in legend on right. A. The 1^st^ row : a conceptual model regarding cells as incompressible spheres with equal radius. B. The 2^nd^ row : cell-arrangement patterns of 222 wild-type embryos. C. The 3^rd^ row : Voronoi segmentation on cells from the close-packing model, using spherical center as origin. D. The 4^th^ row : Voronoi segmentation on cells from the wild-type reference, average position of 222 samples as origin. E. The 5^th^ row : Comparison between close-packing model and wild-type reference. F. The 6^th^ row : Cell-cell contact mapping based on the close-packing model. Intercellular contact was painted with blue, otherwise red or gray (self).

In the conceptual close-packing model, cell-cell contact map is formed deterministically. A series of well-known ground truth involving cell-cell signaling and division-axis regulation can be achieved as following :

● At 4-cell stage, P2 must contact with ABp but not with ABa; Notch signaling : inducing cell-fate differentiation between ABa and ABp^[38–39]^.
● At 4-cell stage and 6-cell stage, P2 must contact with EMS; Wnt signaling : generating two distinct pioneer cells for endoderm (E lineage) and mesoderm (MS lineage)^[18]^.
● At 7-cell stage and 8-cell stage, MS must contact with ABal; Latrophilin signaling : regulating division axis of ABal^[59]^.
● At 8-cell stage, C must contact with ABar; Wnt/*β*-catenin pathway : regulating division axis of ABar^[33]^;

The close-packing configurations in accordance with the wild-type reference can only be rebuilt up to 8-cell stage, presumably for that embryo imaged in our experiment was compressed, which would result in distinct morphogenesis difference from the uncompressed ones since 8-cell stage^[37, 60]^. Besides, cell volume and deformation were ignored here. Even so, the high similarity of structural topology could still support the basic mechanical laws that contribute to cell organization in early *C. elegans* embryo.

## Discussion

Inspired by a fundamental problem on *C. elegans* developmental biology, how could morphogenesis be highly reproducible and precise among individual embryos, this work collected 222 wild-type embryos to establish a reliable statistical reference. After manual editing for quality control and globally linear normalization on space and time, a set of developmental properties in embryo-level as well as single-cell level were obtained quantitatively, including cell cycle, cell division orientation, cell migration trajectory and global growth rate. These wild-type samples together characterized the behaviors and features during embryogenesis, and potentially drive different research topics, such as developmental variability (noise), mutant phenotyping and gene-function inference.

It remains poorly understood that what roles do simple Newtonian mechanics and active molecular regulation play on accurate cell arrangement at early stage. To answer this question, we first analysed the division timing of different cells and identified 8 groups of cells with invariant division order, which provided over 3 minutes on average for system relaxation between any pair of successive division events, while cells within the same group divide synchronously without conserved order. Next, we estimated the spatial cell variability and found that, even though the variability tends to increase over time, the statistical range Δ*r*_STD_ always maintains smaller than cell radius. To figure out if natural coordination on division timing and grouping dominantly contributes to the cell-arrangement accuracy, imaging at time resolution of 10 seconds showed that at the stages with an amount of cell dividing (6-, 12-, 24-cell stages), all the cells would move to the wild-type normal area (Δ*r* < Δ*r*_Q3+1.5*IQR_) in the last 1.5 minutes and keep smaller velocity compared to the earlier one, directly supporting coordinated division timing’s functional impact on system relaxation and cell migration.

However, a few evidences were also found to argue the adequacy of mechanical cues for morphogenesis. Firstly, at some stages with few cells dividing, several cells were found to acquire higher velocity compared to its initial state, seemingly implying that the system could keep developing normally in a motional and noisy embryo environment without relaxation and stabilizing. Secondly, at some extreme wild-type cases, duration between consecutive division events could be quite short but didn’t lead to permanent embryonic defect, meanwhile, cells may be still moving directionally when other cells start to divide, both observations indicate the strong system robustness against intrinsic motional noise. Apart from the wild-type, system-level defect screening on 1818 RNAi-perturbed mutant embryos (758 genes) revealed that no inevitable causality between defect of division timing, division orientation and cell arrangement were found. Note that in many cases the cell-arrangement or division-orientation pattern would spontaneously goes wrong, and sometimes could recover or raise no abnormality followed. These phenomenons suggested that numerous genes and proteins coordinate the morphogenesis in different aspects, along with the driving of mechanical cues. To some extent, underlying molecular regulation provides robustness against perturbation and ability of defect recovery.

Last but not least, a system-level quantitative coding method was designed to represent the phenotype of mutant, by three independent state properties, division timing *T_i_*, division orientation *O_i_* and cell arrangement *A_i_*. The rule could help classify remarkable phenomenons from the mutant database, such as embryo response to perturbation. A Matlab software was further built for looking over our wild-type database, mutant phenotypes and automatically analysing new embryo inputted.

After millions of natural evolution and selection, early *C. elegans* embryo acquired a set of efficient strategies to achieve robust and stable morphogenesis. To balance the trade-off of proliferation rate, cell-positioning robustness and differentiation, combination of simultaneous divisions and ordered divisions was evolved and maintained in early embryogenesis of nematode. Even though distinct division orders provide time for system relaxation and the early cell-arrangement topology seems to accord with simple close-packing model, active molecular regulations from numerous genes and proteins are also critical for morphogenesis. This work provides a statistical wild-type morphogenesis reference and a system-level quantitative dataset of mutant phenotype, and also built a graphical user interface (GUI) using Matlab (MathWorks) for other researchers to look over our wild-type reference, mutant phenotypes and automatically analysing new embryos inputted.

## Acknowledgments

We thank the members of Tang Lab and Zhao Lab for helpful discussion and comments, in particular Prof. Xiaojing Yang and Jingxiang Shen’s constructive advice. The strains were provided by the *C. elegans* Genetic Center (CGC), which is funded by National Institutes of Health, Office of Research Infrastructure Programs, Grant P40 OD010440. This work was supported by the Ministry of Science and Technology of China (2015CB910300), the National Natural Science Foundation of China (91430217), the Hong Kong Research Grants Council (HKBU12100118, HKBU12100917, HKBU12123716) and the HKBU Interdisciplinary Research Cluster Fund. Computation was performed in part on High-Performance Computing Platform at Peking University.

## Author Contributions

C.T., Z.Z., L.T. conceived and coordinated the study. G.G. processed and analysed the data; M.W., V.W.S.H., X.A., L.C. performed RNAi, imaging and embryo curation; B.T. assisted with data computation. C.T., Z.Z., L.T., Z.L. provided theoretical instructions. Z.Z. provided reagents and experimental methods for the project. All the authors reviewed the results and approved of the preprint version of the manuscript.

## Data and Materials Availability

All the relevant data are included within the main paper or the supplementary materials.

## Supplement

The supplement contains supplemental Figures S1-S16, Tables S1-S6 and supplementary materials 1-6 as below :

### Supplementary Material 1

Raw and normalized data of the 222 wild-type embryo samples including division timing from 4- to 350-cell stage and cell position from 4- to 24-cell stage.

### Supplementary Material 2

3D time-lapse image data (time resolution of 1.41 minutes; spatial resolution of 0.09 μm/pixel in focal plane, 0.42 μm/pixel along shooting direction; 60 time points in total) of a wild-type embryo starting from 2-cell stage, which expressed GFP in nucleus (green) and PH(PLC1d1) in membrane (red) (Fig.1B).

### Supplementary Material 3

3D time-lapse image data (time resolution of 10 seconds; spatial resolution of 0.09 μm/pixel in focal plane, 0.42 μm/pixel along shooting direction; 300 time points in total) of a wild-type embryo from 4- to 24-cell stage, which expressed GFP in nucleus (green) (Fig.2DE).

### Supplementary Material 4

Raw and normalized data of the 1818 mutant embryo samples including division timing from 4- to 350-cell stage and cell position from 4- to 24-cell stage.

### Supplementary Material 5

Phenotypic code of reproducible defect after 758 mutant genes perturbed respectively, and particular defect in each mutant embryos (1818).

### Supplementary Material 6

User guide book of Matlab-based software ***STAR 1.0***

---

**Figure S1.**
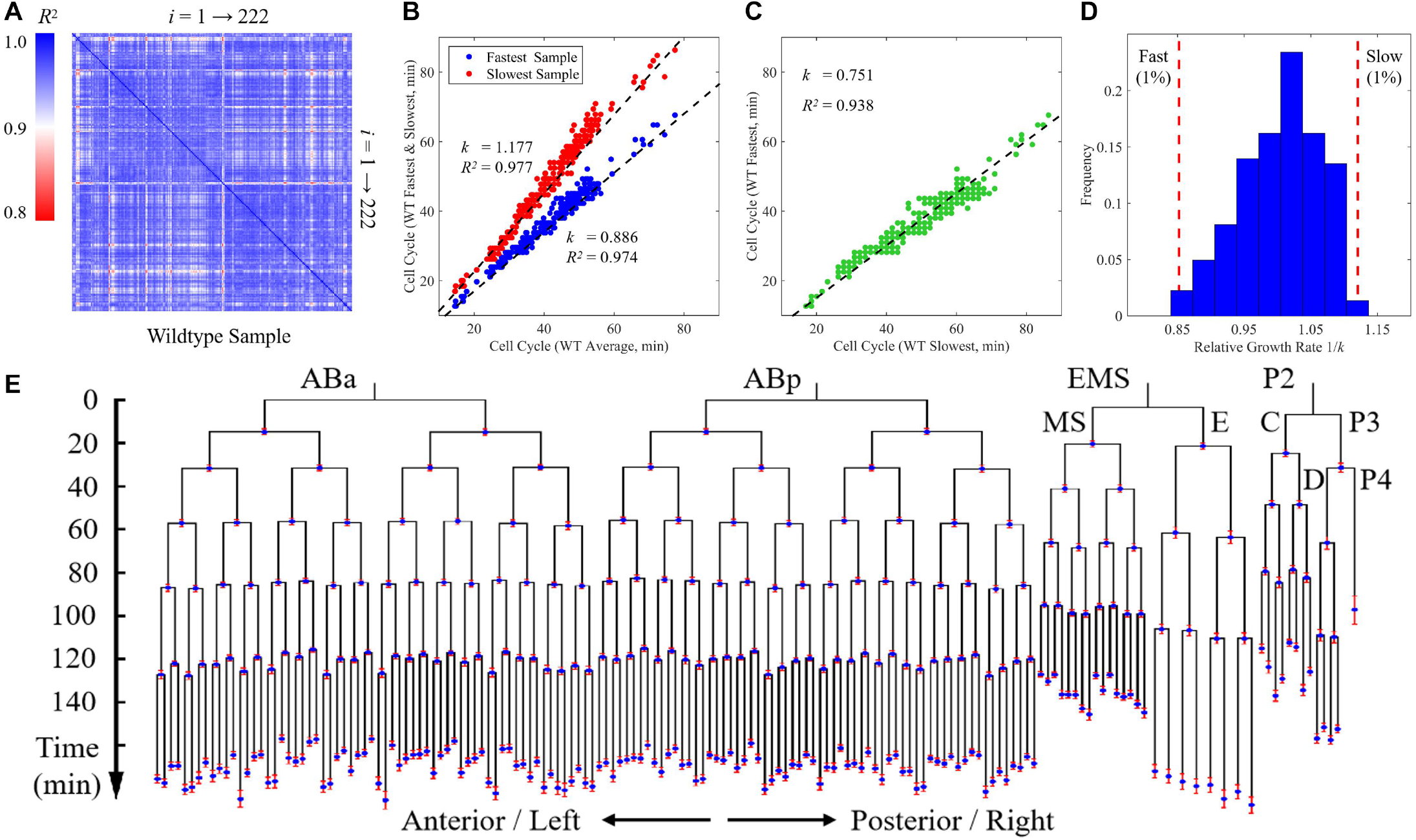
Temporal Information of the 222 wild-type embryos. A. Goodness of fit *R*^2^ of cell cycle between any two wild-type embryos, using proportional function for linear fitting. *i* represents serial number of the 222 wild-type samples (Table S1). Colorbar represents the goodness of fit *R*^2^, ranging from 0.8 to 1.0. B. Comparison between the fastest and slowest embryo samples and the averages of the 222 wild-type embryos. Each point represents a cell with complete lifespan and specific identity, blue for the fastest and red for the slowest embryo; *k*, slope between wild-type averages and the extreme cases; *R*^2^, goodness of fit under proportional function. C. Comparison between the fastest and slowest embryo samples. Each green point represents a cell with complete lifespan and specific identity, plotted with cell cycle in the slowest sample as lateral axis and the one in the fastest sample as vertical axis. D. Histogram of relative global growth rate 1/*k*, with threshold of 1% as extreme outliers illustrated using red dashed lines. E. Cell-lineage tree of *C. elegans* early development consisting of all the cells with complete lifespan (AB4-128, MS1-16, E1-8, C1-8, D1-4, P3 and P4), using average value of normalized cell cycle. Red error bar denotes standard deviation of normalized cell cycle.

**Figure S2.**
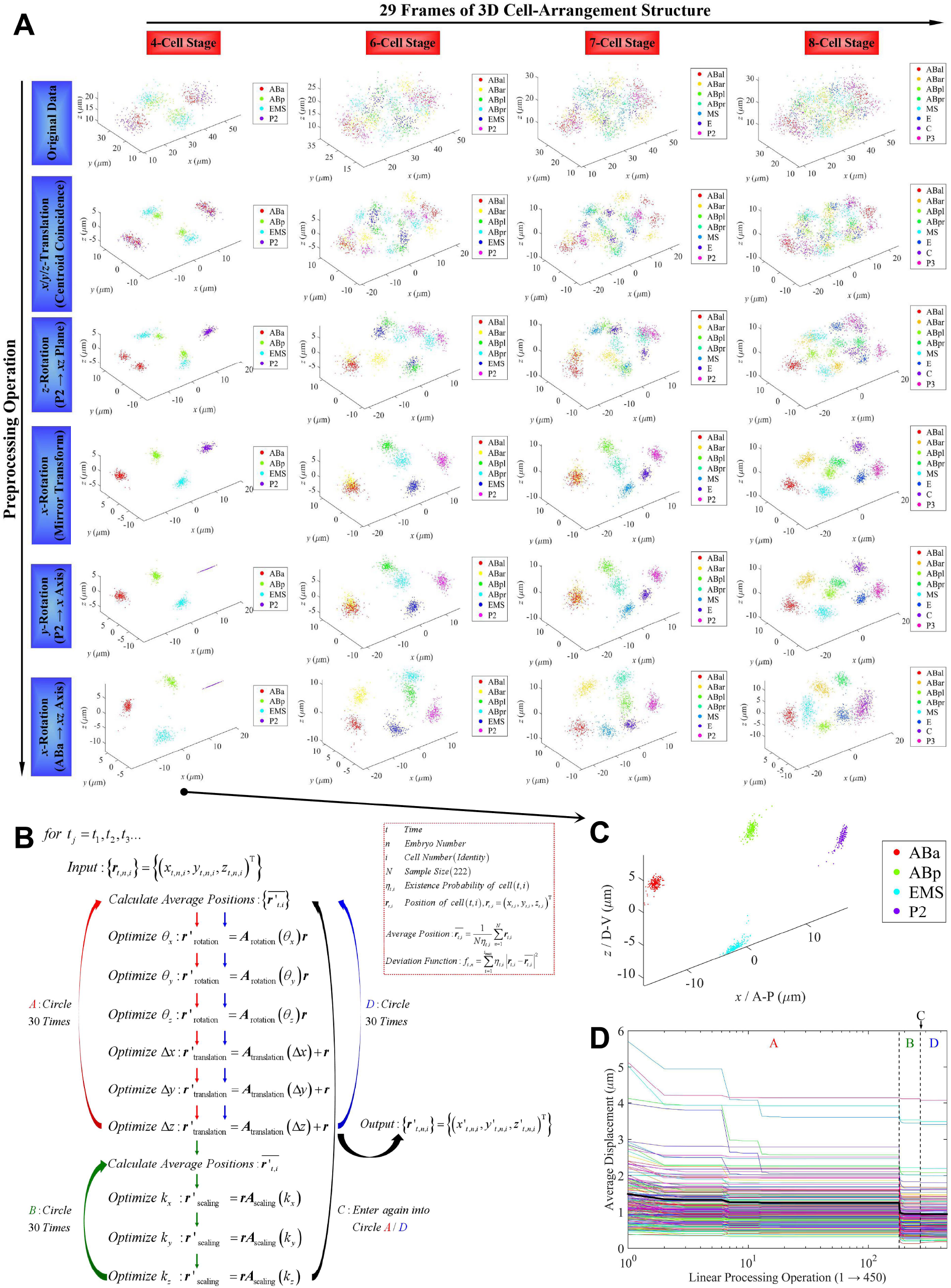
Globally linear normalization on cell positions. A. Linear preprocessing operations for aligning the embryos to rectangular coordinate system, using the multicellular cell structure at 4-cell stage. B. Linear normalization operations for eliminating system variation between the 222 wild-type embryos, using rotation, translation and scaling successively and circularly (Circle A → Circle B → Circle D). C. Normalization result of embryos at 4-cell stage, showing much more concentrated spatial distribution in each cell. D. Average displacement to average cell positions of the 222 wild-type embryos during normalization (450 steps in total). Lines with different color denotes different embryo samples.

**Figure S3.**
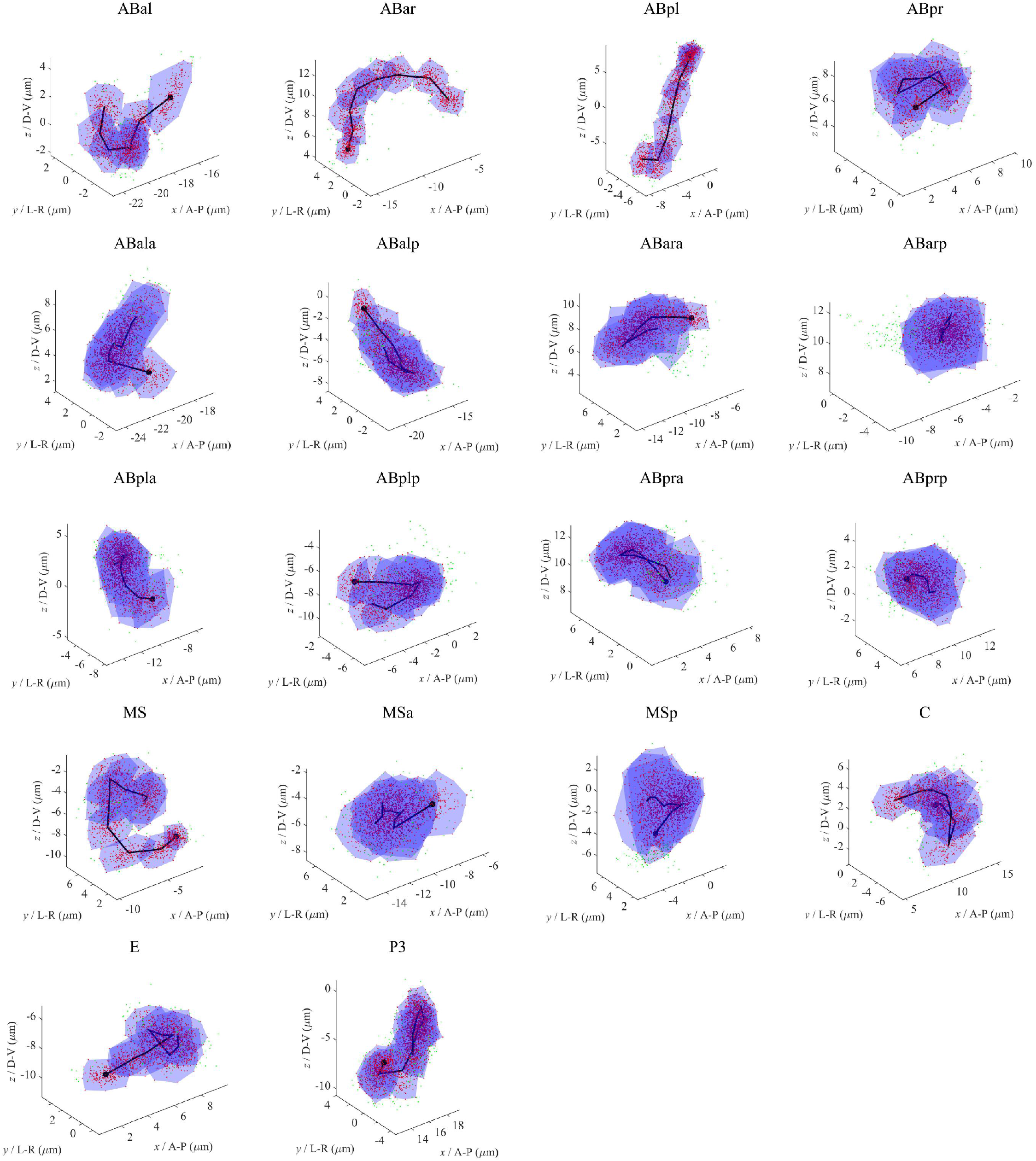
Migration trajectory of cells with complete lifespan during 4- to 24-cell stage. Red points are 95% normal samples forming a convex polyhedron with blue shade, while the other 5% red points are outliers; black line denotes the average migration trajectory, with a black point labeling the origin.

**Figure S4.**
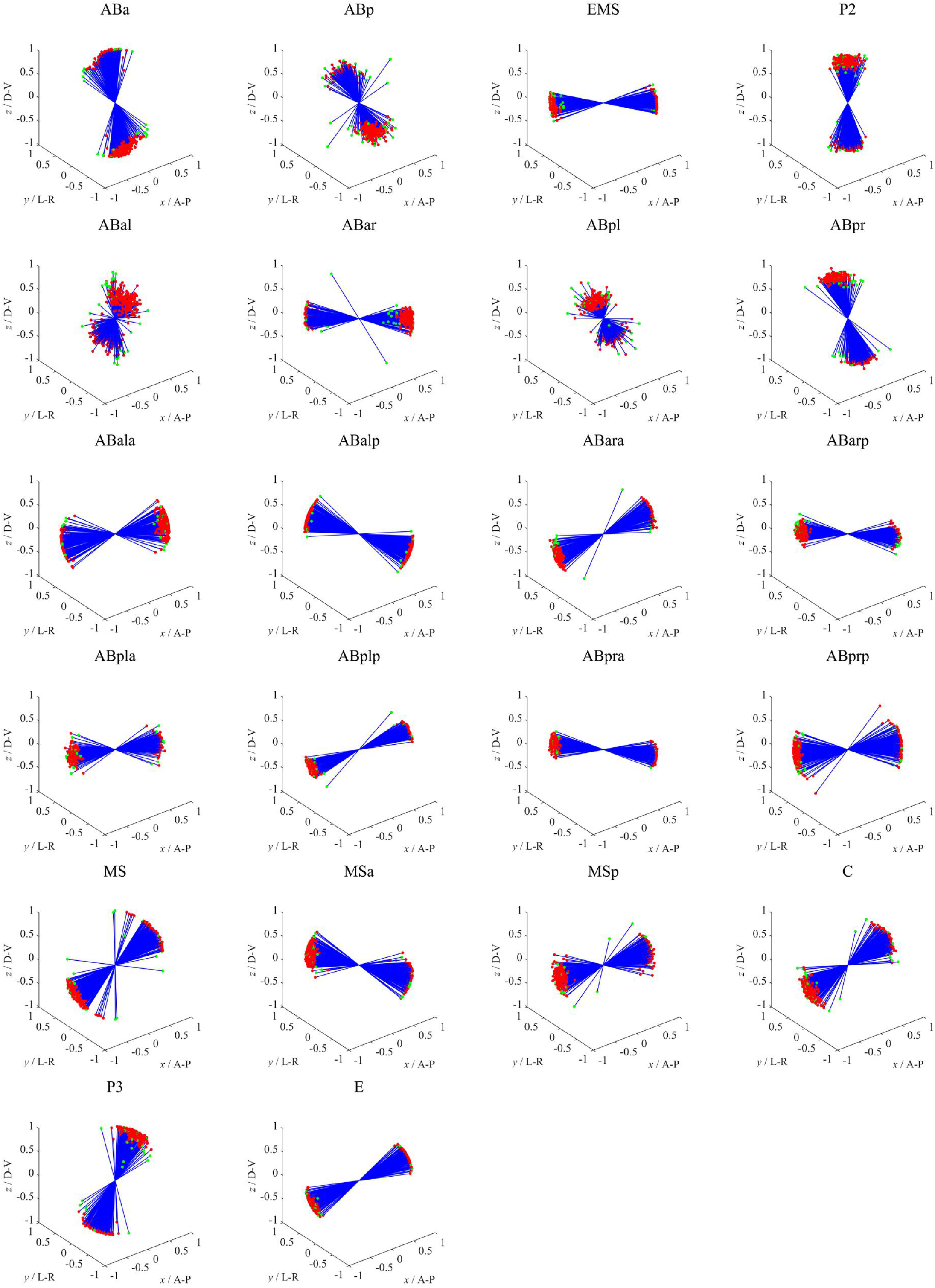
Division-orientation distribution of cells dividing during 4- to 24-cell stage, with initial distance between two daughter cells normalized to one. Red points are 95% normal samples while the other 5% ourliers are painted green.

**Figure S5.**
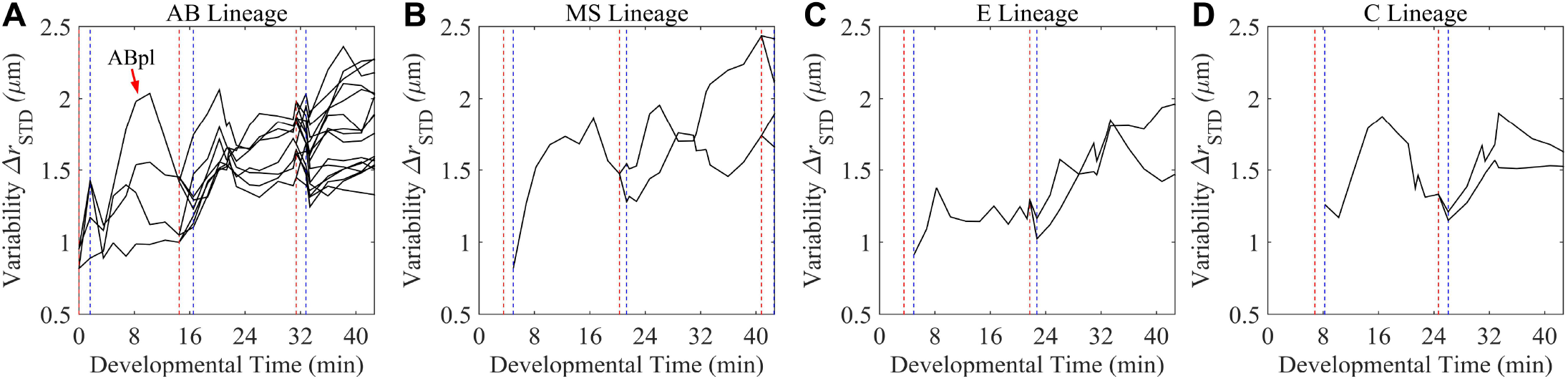
Spatial variability of cells in different lineages reveals fluctuation along with division events. Red and blue dashed lines represent timing of a cell’s last moment and its daughters’ first moment, respectively; black line represents a cell’s spatial variability Δ*r*_STD_. A. AB lineage. Divisions of AB2, AB4 and AB8 are labeled, and the curve of ABpl is highlighted for its highest variability. B. MS lineage. Divisions of EMS, MS and MS2 are labeled. C. E lineage. Divisions of EMS and E are label. D. C lineage. Divisions of P3 and C are labeled.

**Figure S6.**
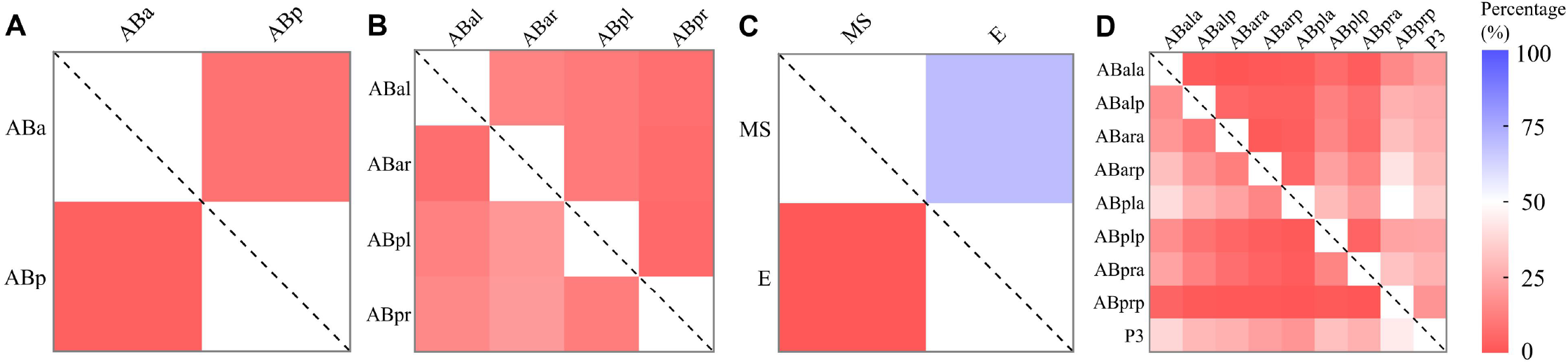
Variation of division timing and ordering of cells within the 4 clustered synchronous cell groups (Table S2). Different colors (i.e. darkness) denote percentage of that a cell on the horizontal axis divides later than other one on the vertical axis (approximately 1.5 minutes at least), using 222 wild-type samples for statistics. No square is painted with the darkest blue, indicating no invariant distinct order exists between any two cells within a group. A. Comparison of division timing between ABa and ABp. B. Comparison of division timing between ABal, ABar, ABpl and ABpr. C. Comparison of division timing between MS and E. In 70.7% of wild-type samples, MS divides at least 1.5 minutes earlier than E, while the others divide within an interval shorter than 1.5 minutes; No embryo obtains an E cell dividing sooner than MS in our wild-type database. D. Comparison of division timing between ABala, ABalp, ABara, ABarp, ABpla, ABplp, ABpra, ABprp, P3.

**Figure S7.**
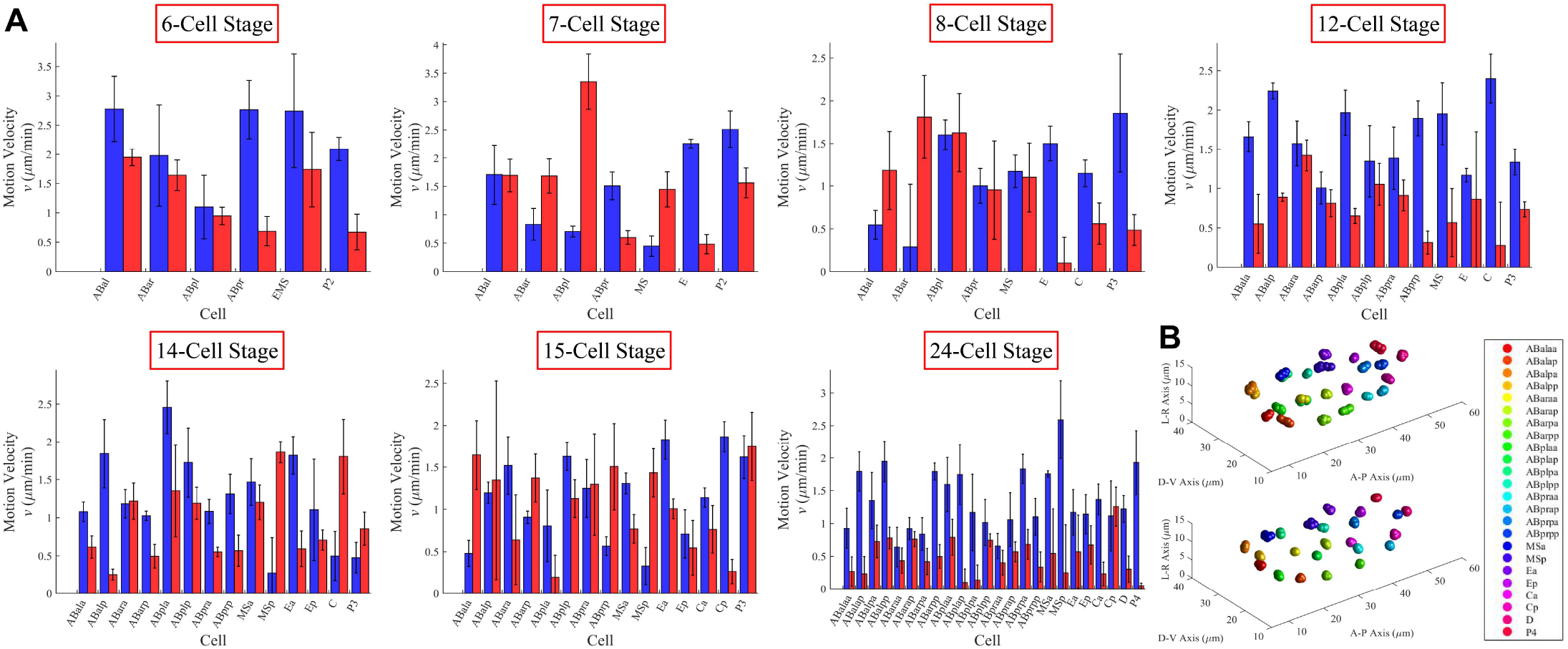
Cell motion in the first and last 1.5 minutes of each stage. A. Cell motion velocity in the first and last 1.5 minutes of each stage. Blue column denotes velocity in the first 1.5 minute; red column denotes velocity in the last 1.5 minutes; error bar denotes standard deviation of velocity in the mean direction. B. Cell motion trajectory in the first (upper figure) and last (lower figure) 1.5 minutes of the 24-cell stage. Each color represents one specific cell identity, noted in legend on right.

**Figure S8.**
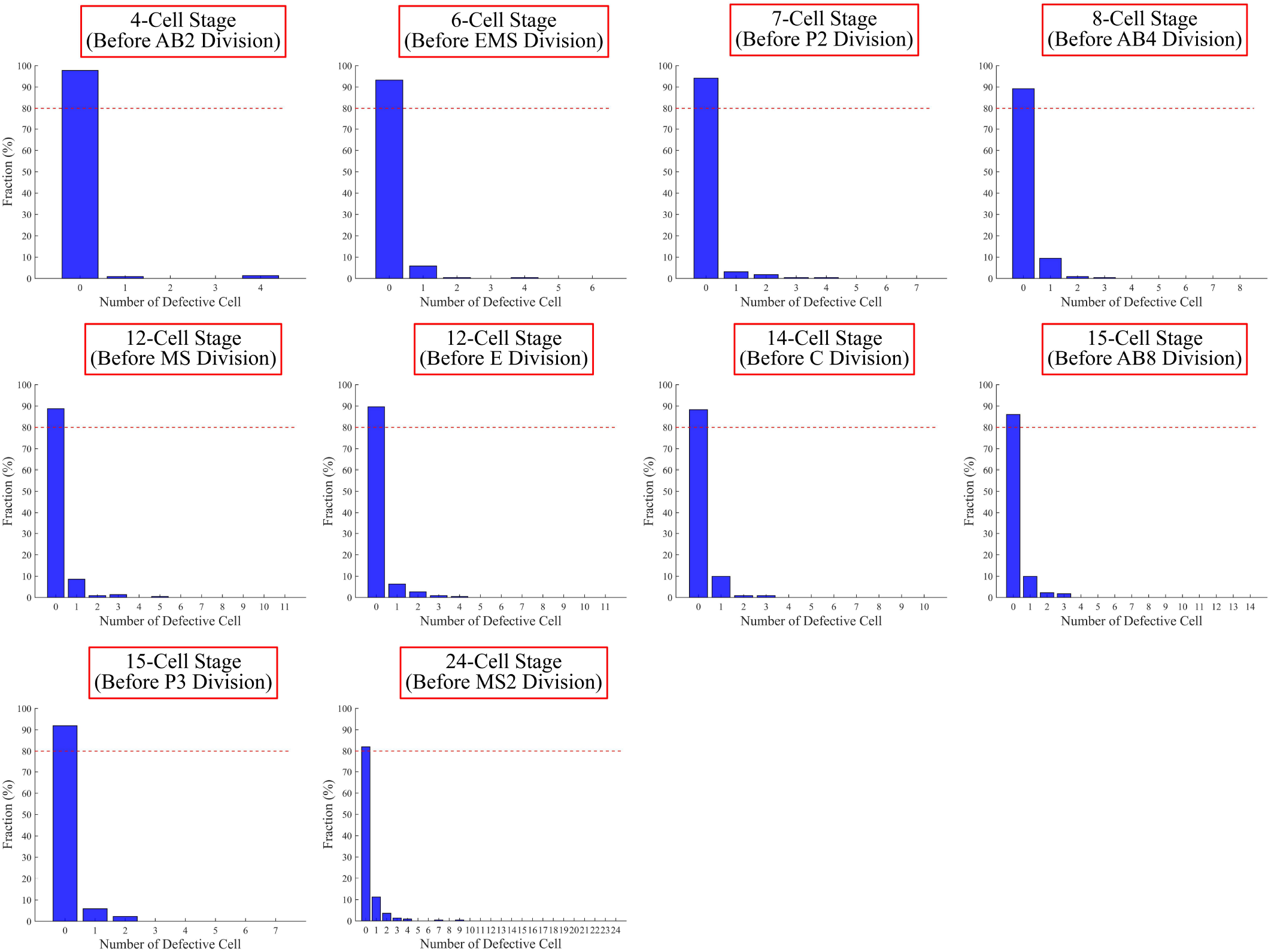
False-positive test on the 12 stages (frames) before cell division events, using cell positions of the 222 wild-type embryos. Blue column denotes the fraction of samples with particular number of defective cells; Over 80% of wild-type embryo samples completely pass through the screening test; for each cell with specific identity, less than 1% of samples would be regarded as defective outliers.

**Figure S9.**
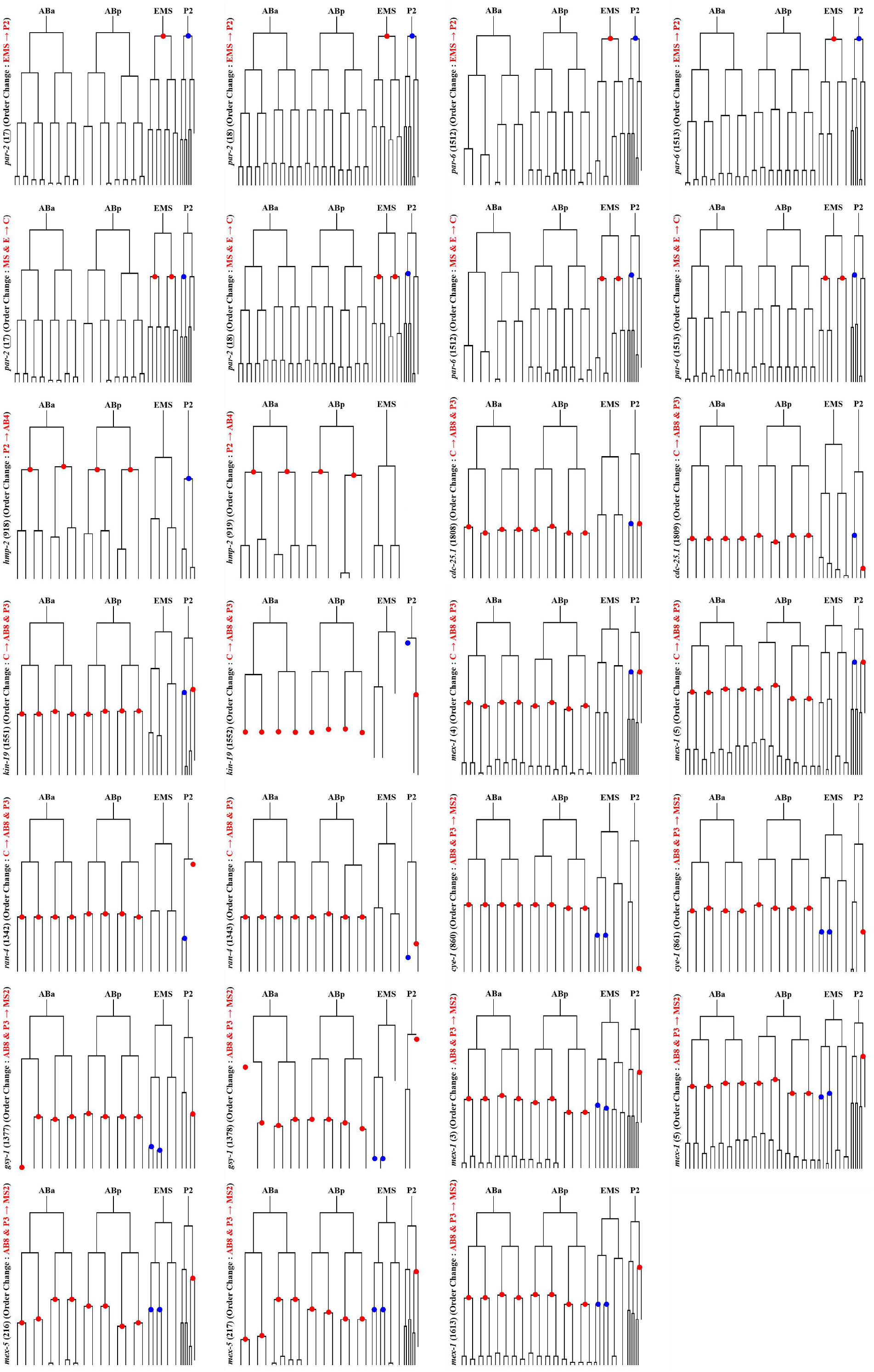
Cell-lineage tree of mutant embryo with reproducible change of division order. Red and blue points denote the two consecutive cell groups with order perturbed (Fig.1C, Table S4); the gene name, serial number of mutant embryo and the natural order are denoted on left; cell without final division time point recorded would not be plotted with black line here.

**Figure S10.**
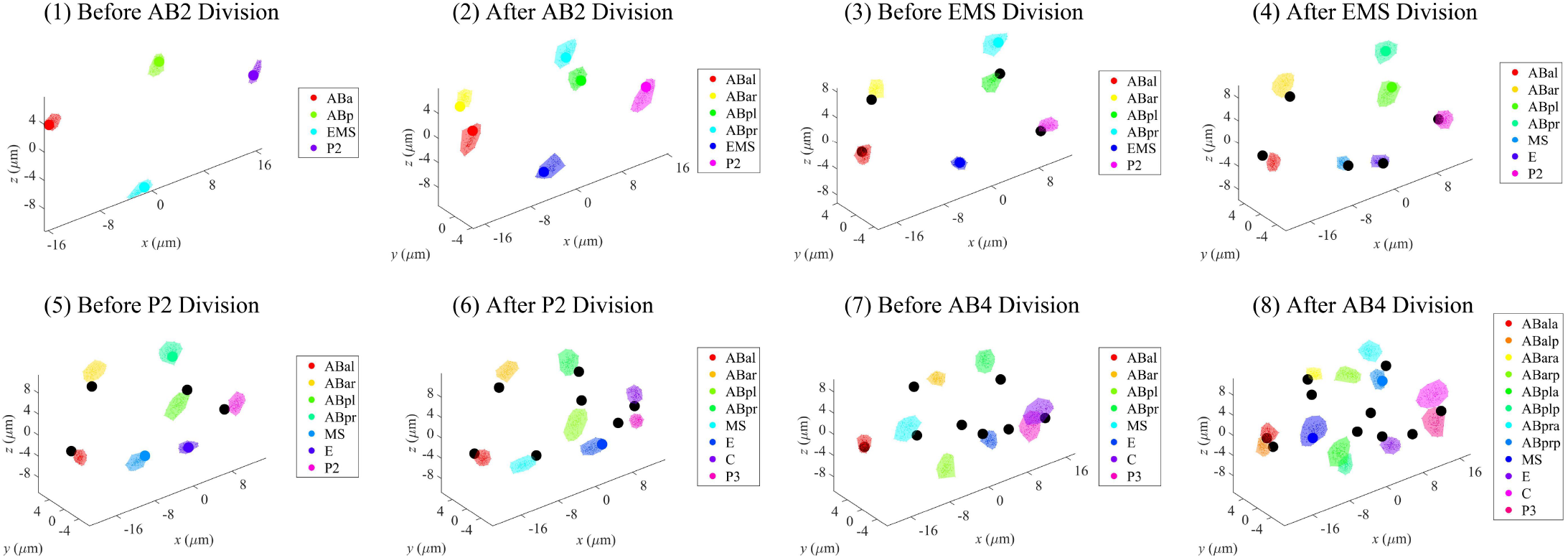
Cell-arrangement and division-orientation progression of *rps-9* RNAi-perturbed embryo (serial number 398, Table S6) during 4- to 12-cell stage, revealing spontaneous cell-arrangement defect without division-timing or division-orientation perturbation. Each color represents one specific cell identity, noted in legend on right; misarranged cells in mutant are illustrated with black point while the normal ones with original color according to the legend.

**Table S1.**
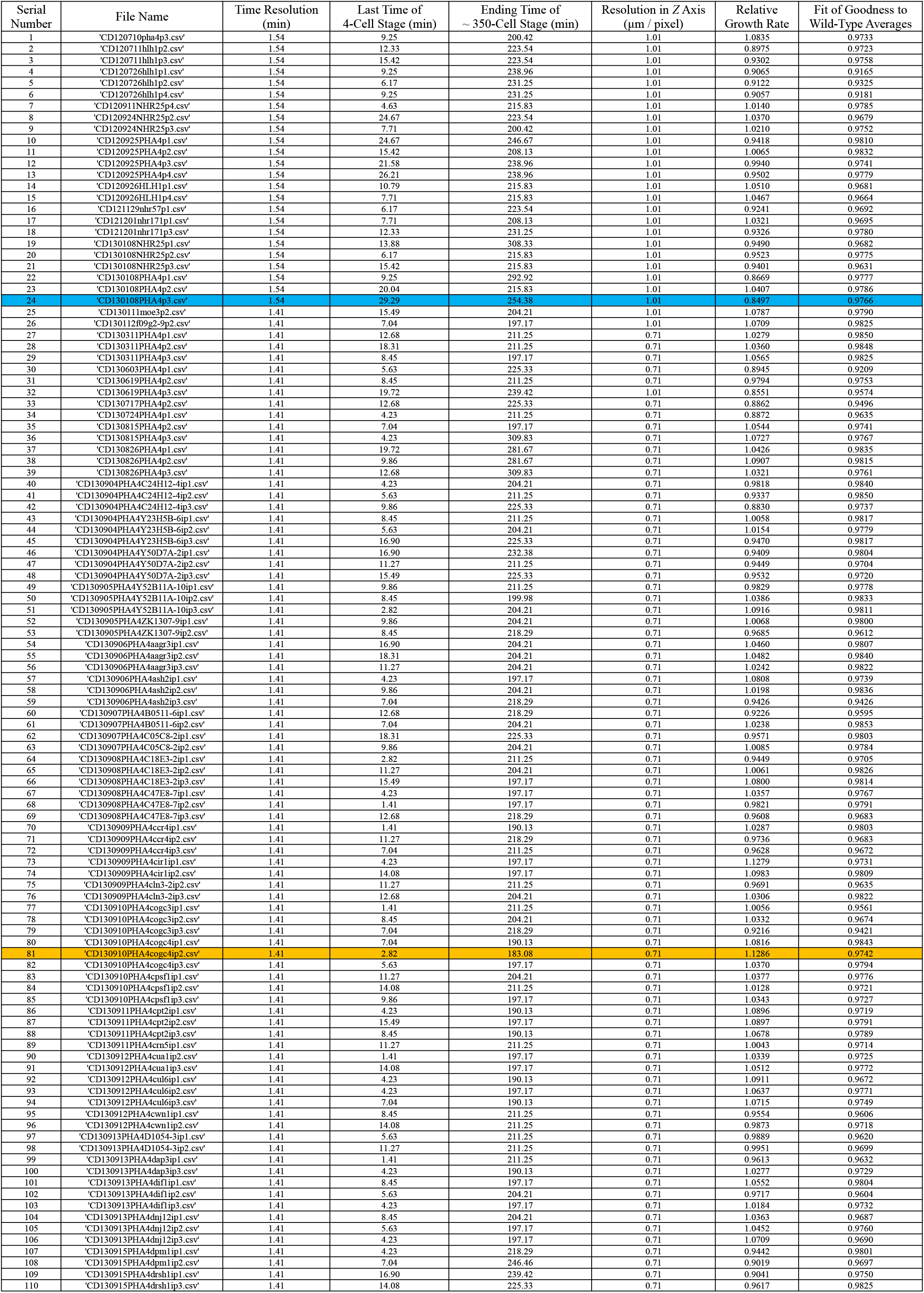
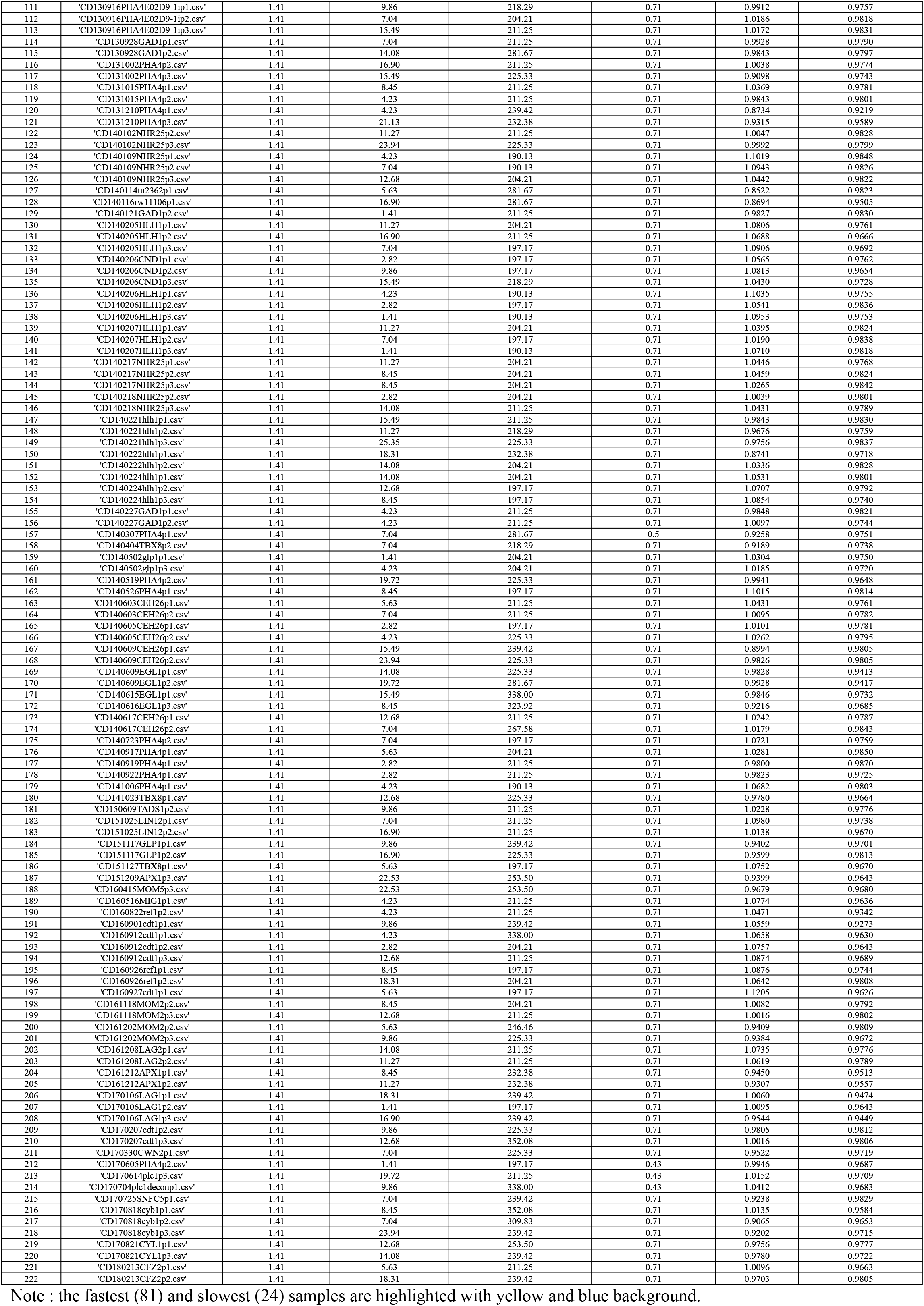
Experimental information about the 222 wild-type embryos.

**Table S2.** Temporal segmentation on developmental process from 4- to 24-cell stage.

| Division Event | First Moment (min) | Last Moment (min) | Inserted Event | Inserted Time (min) |
| --- | --- | --- | --- | --- |
| AB2 | / | 0.00 ± 0.00 [1] | Inserted Event 1 ([8]~[11]) | 10.25 ± 1.33 [9] |
| EMS | / | 3.54 ± 0.91 [3] | Inserted Event 2 ([8]~[11]) | 12.47 ± 1.26 [10] |
| P2 | / | 6.81 ± 1.20 [6] | Inserted Event 3 ([12]~[14]) | 18.63 ± 1.83 [13] |
| AB4 | 1.63 ± 0.57 [2] | 14.49 ± 1.50 [11] | Inserted Event 4 ([19]~[21]) | 20.80 ± 2.56 [20] |
| MS | 4.96 ± 0.92 [4] | 20.32 ± 1.96 [14] | Inserted Event 5 ([25]~[28]) | 36.00 ± 2.94 [26] |
| E | 4.96 ± 0.92 [5] | 21.34 ± 2.04 [15] | Inserted Event 6 ([25]~[28]) | 38.25 ± 3.15 [27] |
| C | 8.23 ± 1.21 [7] | 24.67 ± 2.50 [18] | / | / |
| AB8 | 16.55 ± 1.51 [12] | 30.98 ± 2.62 [21] | / | / |
| P3 | 8.23 ± 1.21 [8] | 31.41 ± 3.04 [22] | / | / |
| MS2 | 21.74 ± 1.97 [16] | 40.86 ± 3.25 [28] | / | / |
| C2 | 26.09 ± 2.51 [19] | / | / | / |
| AB16 | 30.40 ± 2.91 [25] | / | / | / |
| E2 | 22.77 ± 2.06 [17] | / | / | / |
| D | 32.83 ± 3.06 [23] | / | / | / |
| MS4 | 42.68 ± 3.31 [29] | / | / | / |
| P4 | 32.83 ± 3.06 [24] | / | / | / |

Note : Number in blue square bracket represents the order of the 29 events (frames).

**Table S3.** Statistical variation of cell division orientation from experiment.

|  | Cell Name | Angle Variation (Maximum) | Angle Variation (99%) | Angle Variation (95%) |
| --- | --- | --- | --- | --- |
| 1 | ABa | 37.95° | 33.07° | 26.93° |
| 2 | ABp | 45.83° | 28.19° | 21.20° |
| 3 | EMS | 20.27° | 19.62° | 16.69° |
| 4 | P2 | 24.97° | 21.92° | 16.77° |
| 5 | ABal | 34.63° | 30.29° | 23.95° |
| 6 | ABar | 46.25° | 30.32° | 25.10° |
| 7 | ABpl | 32.07° | 26.55° | 19.01° |
| 8 | ABpr | 35.58° | 24.92° | 17.69° |
| 9 | MS | 55.28° | 45.45° | 19.76° |
| 10 | E | 21.04° | 18.61° | 14.99° |
| 11 | C | 44.28° | 26.36° | 19.45° |
| 12 | ABala | 32.21° | 28.13° | 23.68° |
| 13 | ABalp | 26.99° | 24.23° | 20.97° |
| 14 | ABara | 29.90° | 20.78° | 17.48° |
| 15 | ABarp | 19.77° | 17.67° | 14.19° |
| 16 | ABpla | 23.70° | 19.88° | 13.56° |
| 17 | ABplp | 21.12° | 18.74° | 14.13° |
| 18 | ABpra | 17.96° | 16.05° | 13.68° |
| 19 | ABprp | 38.00° | 28.31° | 22.06° |
| 20 | P3 | 42.46° | 36.50° | 24.02° |
| 21 | MSa | 37.41° | 28.08° | 21.07° |
| 22 | MSp | 49.51° | 24.43° | 18.39° |

Note : To quantify the division-angle variation, the narrowest cone encircling the arbitrary percentage (100%, 99%, 95%) of all the experimental division vectors was searched and identified.

**Table S4A.** Invariant division order during 4- to 24-cell stage in wild-type embryo.

|  | Division Order |  |  | Time Interval (min)<br>on Average | Minimum (min) | Maximum (min) | Percentage |
| --- | --- | --- | --- | --- | --- | --- | --- |
| 1 | AB2* | → | EMS | 3.38 ± 0.98 | 1.23 | 8.83 | 100% |
| 2 | EMS | → | P2 | 3.27 ± 0.82 | 1.38 | 6.98 | 100% |
| 3 | P2 | → | AB4* | 7.69 ± 0.93 | 4.92 | 10.4 | 100% |
| 4 | AB4* | → | MS&E* | 5.12 ± 1.12 | 1.47 | 8.85 | 100% |
| 5 | MS&E* | → | C | 3.35 ± 1.04 | 1.33 | 8.39 | 100% |
| 6 | C | → | AB8&P3* | 6.07 ± 1.04 | 1.52 | 8.53 | 100% |
| 7 | AB8&P3* | → | MS2* | 10.14 ± 1.27 | 7.03 | 13.84 | 100% |

\* denotes a group with more than one cells.

**Table S4B.**
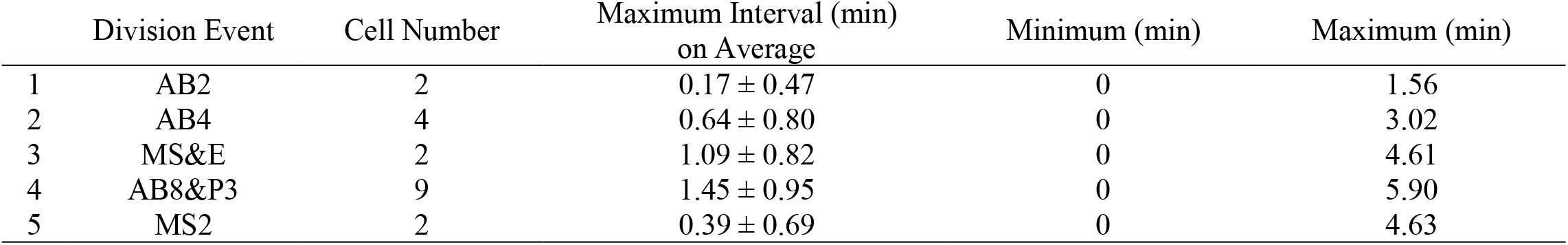
Maximum interval between division timing of cells within the same group.

**Table S5.**
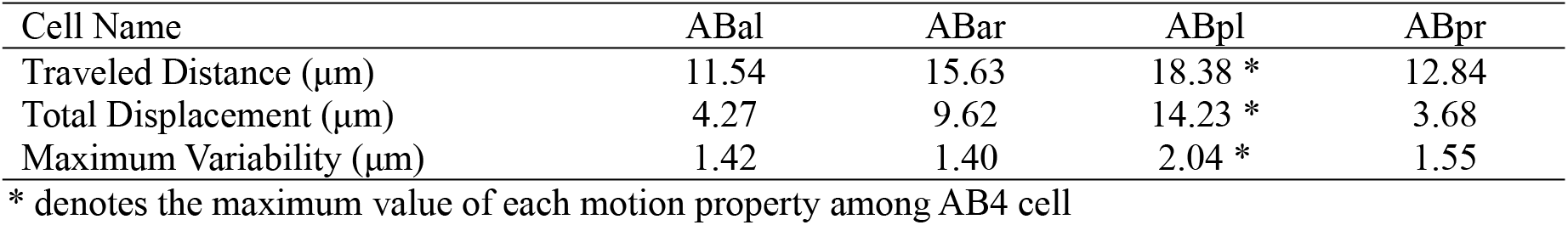
Comparison of motion properties between AB4 cells.

| Cell Name | ABal | ABar | ABpl | ABpr |
| --- | --- | --- | --- | --- |
| Traveled Distance (μm) | 11.54 | 15.63 | 18.38 * | 12.84 |
| Total Displacement (μm) | 4.27 | 9.62 | 14.23 * | 3.68 |
| Maximum Variability (μm) | 1.42 | 1.40 | 2.04 * | 1.55 |

\* denotes the maximum value of each motion property among AB4 cell

**Table S6.**
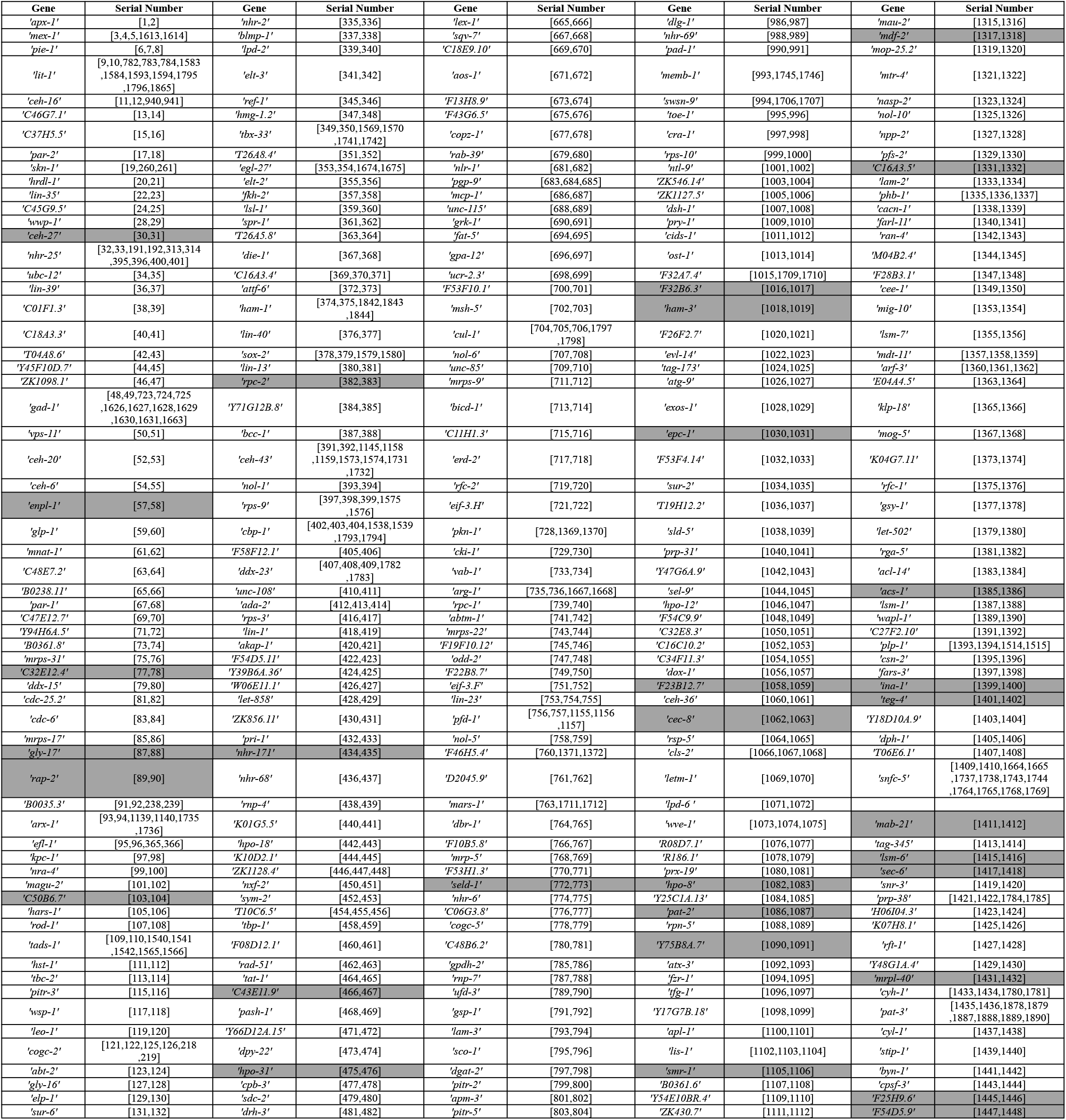
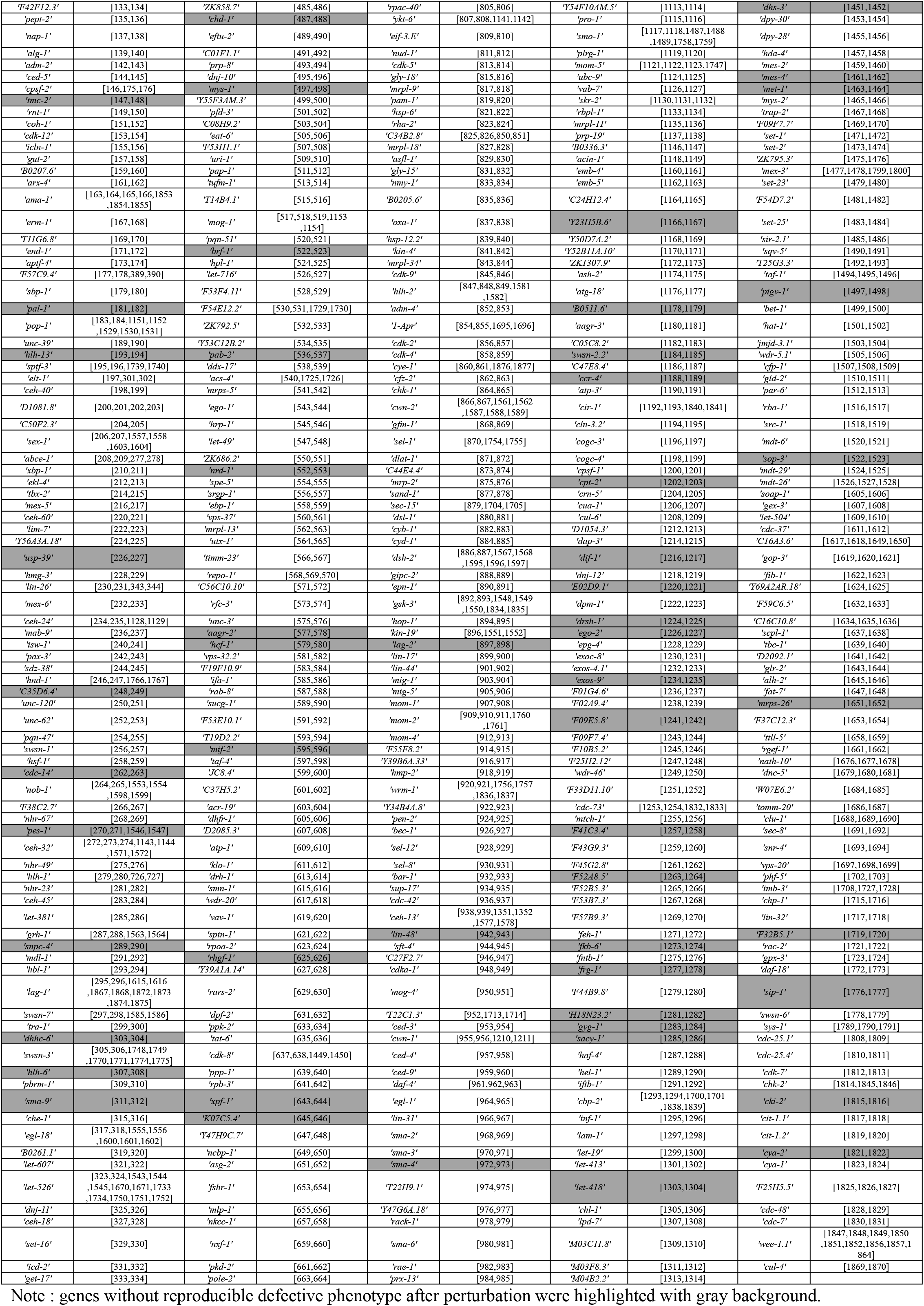
Information about the 758 RNAi-perturbed genes.

## Supplementary Material 6. User guide book of software *STAR 1.0*

### **1.** Summary

● ***STAR 1.0*** (**<u>S</u>**patio-**<u>T</u>**emporal **<u>A</u>**tlas at single-cell **<u>R</u>**esolution of *C. elegans* morphogenesis) is an integrated software designed mainly for three functions :
■ ***Function 1*** : Looking over the morphogenesis reference formed by 222 wild-type embryo samples;
■ ***Function 2*** : Looking over the mutant phenotypes under perturbation of 758 genes respectively;
■ ***Function 3*** : Automatically phenotyping and analysing new inputted embryos.
● This software was designed with Matlab 2015a and could also be run on the higher versions of Matlab.
● After downloading the whole folder named **GUI-STAR1.0**, 6 subfolders would be acquired as the following : (Fig.S11A)
■ **DataStorage** : Files inside are results after analysis of new inputted embryo.
■ **Example** : A .csv file (*mex-5*, serial number 217) is used to be analysed by ***Function 3*** as a test example.
■ **MutantCode-1** : Files inside are reproducible phenotypes of the 758 mutant, named with their gene names (Fig.S11B).
■ **MutantCode-2** : Files inside are phenotypes of the 1818 mutant embryos, named with their gene names and serial numbers (Table S6, 6Fig.S11C).
■ **MutantDatabase** : 2*1818 files of morphogenesis information of the mutant database.
■ **Tool** : Files inside are data of wild-type reference and tools for computation.
● The .m files and .fig files here are critical subfunctions and components of ***STAR 1.0***, any deletion of them may make the software lose its originally designed functions.
● To initiate ***STAR 1.0***, one needs to use Matlab to open the **STAR.m** file, and click the **Run** button on top. (Fig.S11D)
● The ***STAR 1.0*** graphical user interface mainly includes five parts (Fig.S11E) :
■ ***Function 1 (Wild-Type Reference)*** : Both global cell-arrangement pattern and single-cell developmental property (cell cycle, division orientation, migration trajectory) of the statistical wild-type reference can be visualized.
■ ***Function 2 (Mutant Phenotype)*** : Both global cell-arrangement pattern and single-cell developmental property (cell cycle, division orientation, migration trajectory) of the mutant embryos can be visualized along with their analysed results (defective or not).
■ ***Function 3 (Defect Detection)*** : Both global cell-arrangement pattern and single-cell developmental property (cell cycle, division orientation, migration trajectory) of the new inputted embryos can be visualized along with their analysed results (defective or not).
■ ***Program State*** : running state of the software would be shown, such as “Figure Plotting” and “Plotting Finished”.
■ ***Value Assignment*** : According to the segmentation method mentioned (Fig.3D, Table S2), 10 division events are highlighted and labeled with number according to their average division timing as well as order : AB2 (1), EMS(2), P2(3), AB4(4), MS(5), E(6), C(7), P3(8), AB8(9), MS2(10). Besides, moments before and after a division are marked with 1 and 2 respectively. These arbitrarily designed corresponding relationships between division event and serial number assists users to look over their interested developmental stage, which is also noted in red on the user interface.

**Figure S11.**
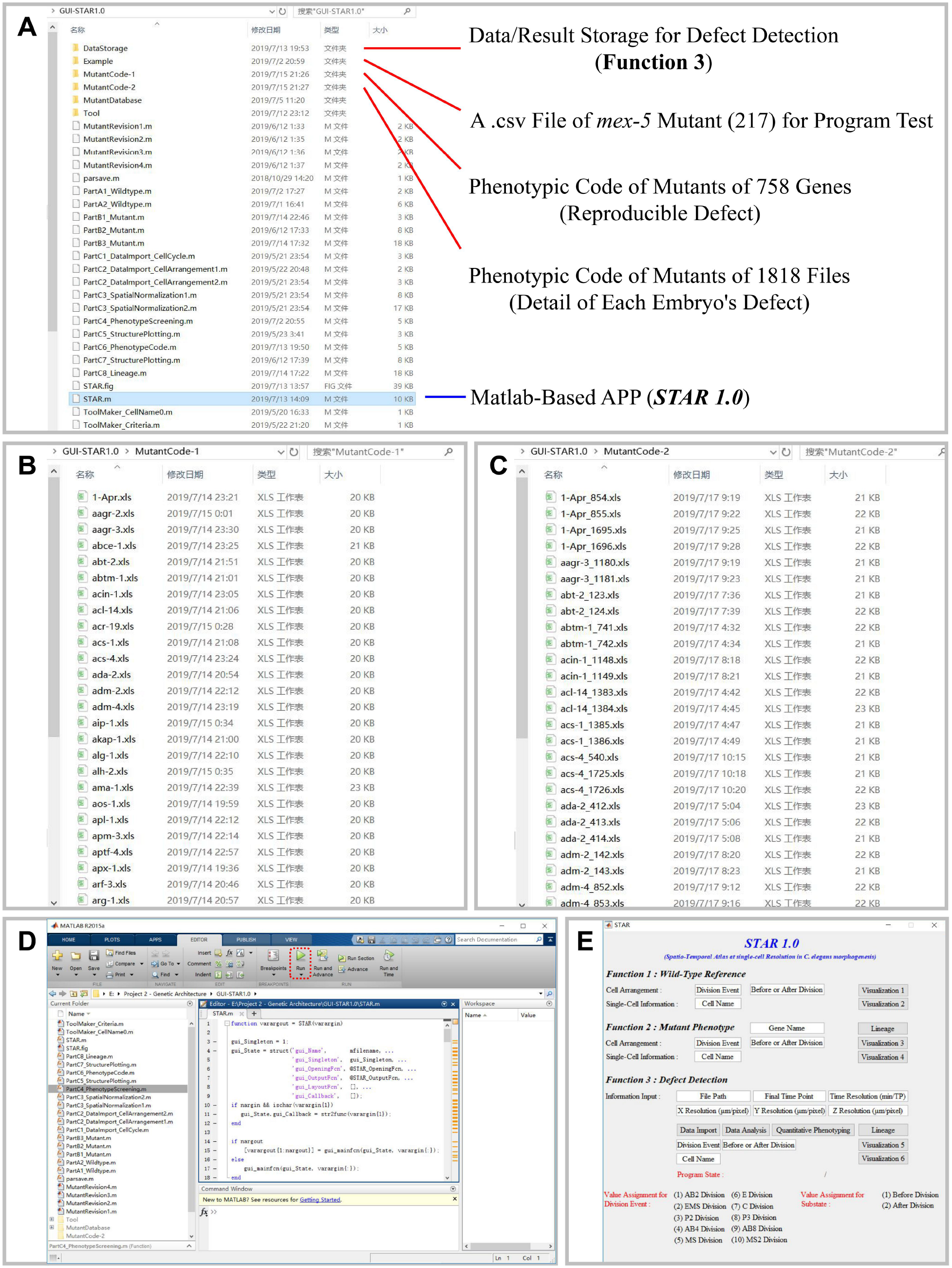
Major components of software ***STAR 1.0***. A. The folder **GUI-STAR1.0** consists of six subfolders and several function documents assisting the program. The **STAR.m** file, namely the graphical user interface, is highlighted with blue shade. B. The 758 files containing reproducible phenotypes of mutant are listed and named with their gene names (details introduced in Fig.S15). C. The 1818 files containing phenotypes of all the mutant embryos are listed and named with their gene names followed by serial numbers (Table S6, details introduced in Fig.S16). D. The Matlab window after opening the **STAR.m** file. The user should click the **Run** button on top to initiate ***STAR 1.0***. E. Graphical user interface of ***STAR1.0***, mainly including five parts : ***Function 1 (Wild-Type Reference)***, ***Function 2 (Mutant Phenotype)***, ***Function 3 (Defect Detection)***, ***Program State*** and ***Value Assignment***.

### **2.** Function Introduction

#### ● Function 1 (Wild-Type Reference)

This function enables user to look over the global cell-arrangement pattern and single-cell developmental properties including cell cycle, division orientation and migration trajectory from 4- to 24-cell stage, using the 222 wild-type embryo samples. Please test and use the software following the steps below.

A. Cell-Arrangement Visualization (Fig.S12A)

a. Input the serial number of division event into the “Division Event” text box, referring to the *Value Assignment* part;
b. Input the assigned value of substate (1, before division; 2, after division) into the “Before or After Division” text box;
c. Click the “Visualization 1” button and wait for seconds;
d. The interface will show “Figure Plotting” on the right of the “Program State” text when the program running, and show “Plotting Finished” after output of the figure.
B. Single-Cell Property Visualization (Fig.S12B)

a. Input the name of cell into the “Cell Name” text box;
b. Click the “Visualization 2” button and wait for seconds;
c. The interface will show “Figure Plotting” on the right of the “Program State” text when the program running, and show “Plotting Finished” after out put of the figure.

**Figure S12.**
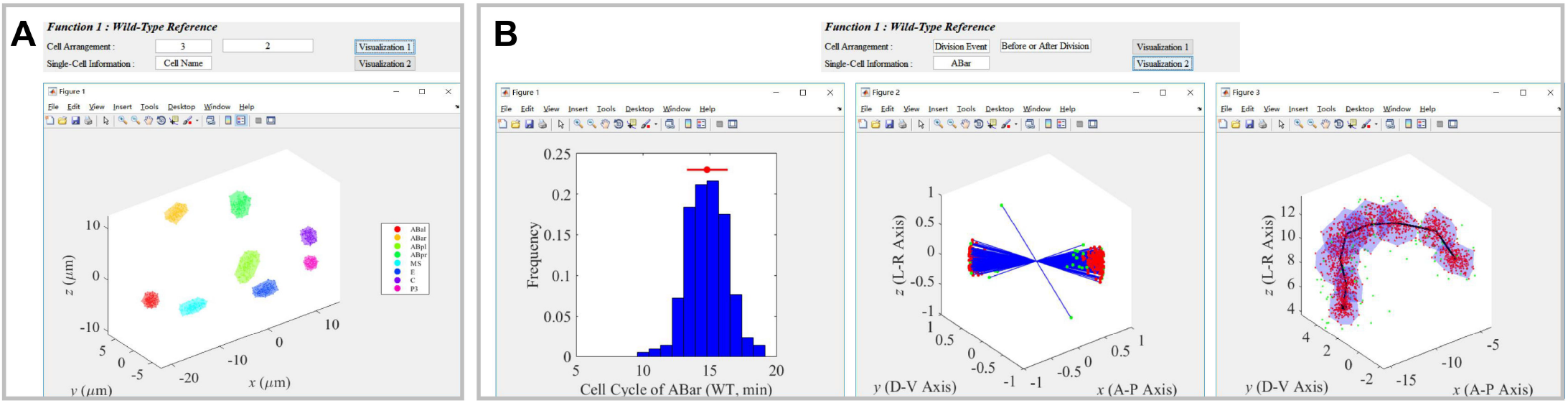
Introduction to ***Function 1 (Wild-Type Reference)***. A. Global cell-arrangement pattern was formed and shown with 222 wild-type samples, after inputting the assigned values of division event and substate interested and clicking the “Visualization 1” button. Convex polyhedron formed by 95% wild-type normal points of each cell is illustrated; each color represents one specific cell identity, noted in legend on right. B. Single-cell developmental properties including cell cycle, division orientation and migration trajectory was formed and shown with 222 wild-type samples, after inputting the identity name of cell interested and clicking the “Visualization 2” button. For cell-cycle distribution, the average and standard deviation are exhibited with red point and line (AVERAGE ± STD) on top. For division-orientation distribution, the initial distances between two daughters were normalized to one; Red points are 95% normal samples while the other 5% are outliers. Migration-trajectory distribution was formed by 95% normal samples in red which form a convex polyhedron with blue shade, and the other 5% as outliers in green; black line denotes the average migration trajectory, with a black point labeling the origin.

#### ● Function 2 (Mutant Phenotype)

This function enables user to look over the global cell-arrangement pattern and single-cell developmental properties including cell cycle, division orientation and migration trajectory from 4- to 24-cell stage, using the 1818 mutant embryo samples. For each gene, all the files belonging to it would be shown (Table S6). Please test and use the software following the steps below.

A. Cell-Lineage Tree Visualization (Fig.S13A)

a. Input the name of gene into the “Gene Name” text box;
b. Click the “Lineage” button and wait for seconds;
c. The interface will show “Figure Plotting” on the right of the “Program State” text when the program running, and show “Plotting Finished” after output of the figure.
B. Cell-Arrangement Visualization (Fig.S13B)

a. Input the name of gene into the “Gene Name” text box;
b. Input the serial number of division event into the “Division Event” text box, referring to the *Value Assignment* part;
c. Input the assigned value of substate (1, before division; 2, after division) into the “Before or After Division” text box;
d. Click the “Visualization 3” button and wait for seconds;
e. The interface will show “Figure Plotting” on the right of the “Program State” text when the program running, and show “Plotting Finished” after out put of the figure.
C. Single-Cell Property Visualization (Fig.S13C)

a. Input the name of cell into the “Cell Name” text box;
b. Click the “Visualization 4” button and wait for seconds;
c. The interface will show “Figure Plotting” on the right of the “Program State” text when the program running, and show “Plotting Finished” after out put of the figure.

**Figure S13.**
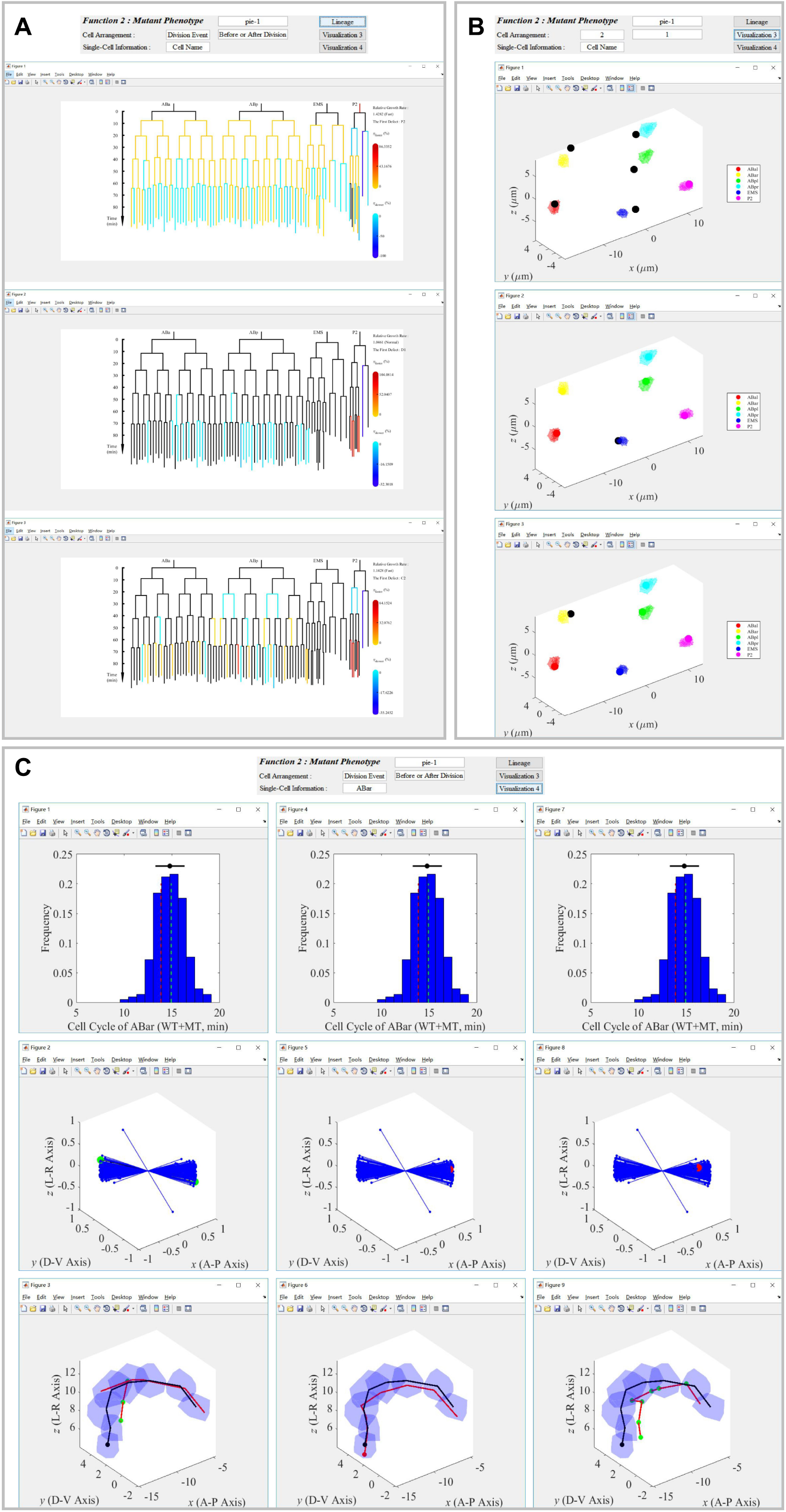
Introduction to ***Function 2 (Mutant Phenotype)***. A. Cell-lineage tree is plotted using cells with complete lifespan in the mutant file as well as AB2, EMS and P2 cells. Cells with normal cell cycle is drawn in black, while the one dividing significantly faster in red and slower in blue; level of cell-cycle defect is quantified with relative deviation *η* = (Mutant Cell Cycle - WT Average) / (WT Average), denoted with colorbar on right; the relative global growth rate and the first division-timing defect are shown at the right top; the developmental time is shown on left with a vertical axis, using the last moment of 4-cell stage as the origin. B. Global cell-arrangement patterns of all the mutant samples belonging to the gene inputted are plotted, after inputting the assigned values of division event and substate interested and clicking the “Visualization 3” button; convex polyhedron formed by 95% wild-type normal points of each cell is illustrated; each color represents one specific cell identity, noted in legend on right; misarranged cell (false positive ≤ 0.01) in mutant is highlighted with black point while the normal one with ordinary color. C. Single-cell developmental properties including cell cycle, division orientation and migration trajectory was formed and shown with 222 wild-type samples, after inputting the identity name of cell interested and clicking the “Visualization 2” button. For cell-cycle distribution, the average and standard deviation are exhibited with black point and line (AVERAGE ± STD) on top, while original and normalized cell cycles of mutant are illustrated with dashed line in red and green respectively. For division-orientation distribution, the initial distances between two daughters were normalized to one; all the wild-type samples are shown with blue line and point, while the mutant one was highlighted with larger point in red if normal or in green if defective (false positive ≤ 0.01). For migration-trajectory distribution, convex polyhedron formed by 95% normal wild-type samples were shown with blue shade; black line denotes the average migration trajectory, with a black point labeling the origin; red line denotes mutant migration trajectory; misarranged cell is highlighted with green point, otherwise red.

#### ● Function 3 : Defect Detection

This function provides automatic and quantitative analysis on new embryo inputted, and also enables user to look over the global cell-arrangement pattern and single-cell developmental properties including cell cycle, division orientation and migration trajectory from 4- to 24-cell stage, using the 222 wild-type embryos as reference. Please test and use the software following the steps below. Note that the .csv file (Path : Example\CD121112PHA4mex5ip3.csv) can be used as an example for test. The analysis results will be stored with an .xls file named “PhenotypeCode.xls” in the **DataStorage** subfolder.

A. **Data Preprocessing** (Fig.S14A)

a. Input the file path of csv-format embryo data into the “File Path” text box and wait for seconds;
b. Input the ending time point of valid data into the “Final Time Point” text box;
c. Input the time resolution of imaging into the “Time Resolution (min/TP)” text box;
d. Input the spatial resolution in x direction (confocal plane) into the “X Resolution (μm/pixel)” text box;
e. Input the spatial resolution in y direction (confocal plane) into the “Y Resolution (μm/pixel)” text box;
f. Input the spatial resolution in z direction (shooting direction) into the “Z Resolution (μm/pixel)” text box;
g. Click the “Data Import” button and wait for seconds;
h. The interface will show “Cell-Cycle Data Importing” on the right of the “Program State” text when the program running, and show “Data Import Finished” after output of the figure;
i. Click the “Data Analysis” button and wait for seconds;
j. The interface will show “Data Analysing” on the right of the “Program State” text when the program running, and show “Data Analysis Finished” after output of the figure;
k. Click the “Quantitative Phenotyping” button and wait for seconds;
l. The interface will show “Phenotyping” on the right of the “Program State” text when the program running, and show “Phenotyping Finished” after output of the figure.
B. **Cell-Arrangement Visualization** (Fig.S14B)

a. Input the serial number of division event into the “Division Event” text box, referring to the **Value Assignment** part;
b. Input the assigned value of substate (1, before division; 2, after division) into the “Before or After Division” text box;
c. Click the “Visualization 5” button and wait for seconds;
d. The interface will show “Figure Plotting” on the right of the “Program State” text when the program running, and show “Plotting Finished” after out put of the figure.
C. **Cell-Lineage Tree Visualization** (Fig.S14C)

a. Click the “Lineage” button and wait for seconds;
b. The interface will show “Figure Plotting” on the right of the “Program State” text when the program running, and show “Plotting Finished” after output of the figure;
D. **Single-Cell Property Visualization** (Fig.S14D)

a. Input the name of cell into the “Cell Name” text box.
b. Click the “Visualization 6” button and wait for seconds.
c. The interface will show “Figure Plotting” on the right of the “Program State” text when the program running, and show “Plotting Finished” after out put of the figure.

**Figure S14.**
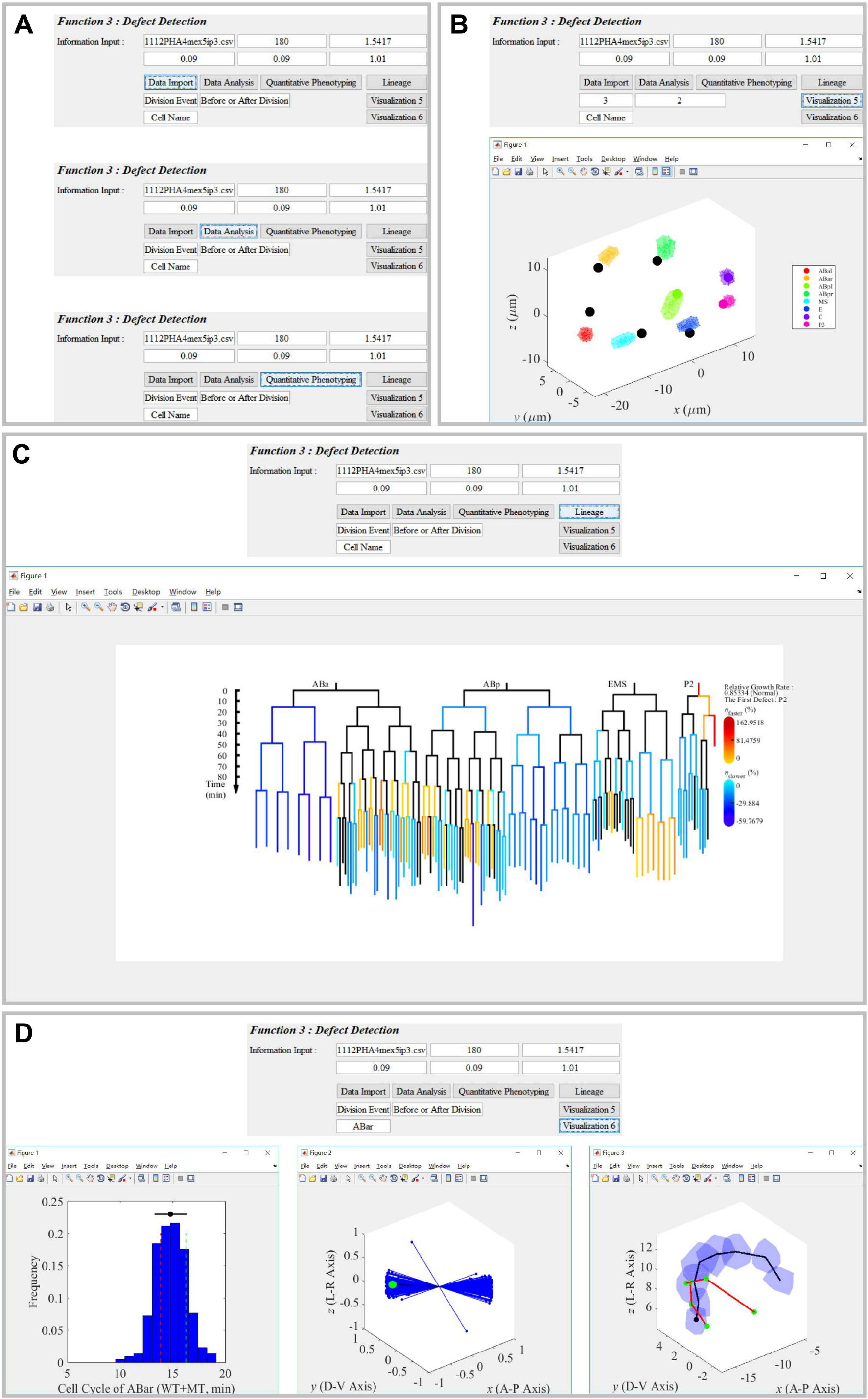
Introduction to ***Function 3 (Defect Detection)***. A csv-format file placed in the **Example** subfolder was used for test and as an example of required data format. A. Preprocessing on the new file requires the user to input the experimental information and click the “Data Import”, “Data Analysis”, “Quantitative Phenotyping” buttons successively. B. Global cell-arrangement pattern is calculated and plotted, after inputting the assigned values of division event and substate interested and clicking the “Visualization 5” button; convex polyhedron formed by 95% wild-type normal points of each cell is illustrated; each color represents one specific cell identity, noted in legend on right; misarranged cell (false positive ≤ 0.01) in the embryo inputted is highlighted with black point while the normal one with ordinary color. C. Cell-lineage tree was plotted using cells with complete lifespan in the embryo inputted as well as AB2, EMS and P2 cells. Cells with normal cell cycle was drawn in black, while the one dividing significantly faster in red and slower in blue; level of cell-cycle defect is quantified with relative deviation *η* = (Mutant Cell Cycle - WT Average) / (WT Average), denoted with colorbar on right; The relative global growth rate and the first division-timing defect are shown at the right top; The developmental time is shown on left with a vertical axis, using the last moment of 4-cell stage as the origin. D. Single-cell developmental properties including cell cycle, division orientation and migration trajectory was formed and shown, after inputting the identity name of cell interested and clicking the “Visualization 6” button. For cell-cycle distribution, the average and standard deviation are exhibited with black point and line (AVERAGE ± STD) on top, while original and normalized cell cycles are illustrated with dashed line in red and green respectively. For division-orientation distribution, the initial distances between two daughters are normalized to one; all the wild-type samples are shown with blue line and point, while the one of new inputted embryo was highlighted with larger point in red if normal or in green if defective (false positive ≤ 0.01). For migration-trajectory distribution, convex polyhedron formed by 95% normal wild-type samples were shown with blue shade; black line denotes the average migration trajectory, with a black point labeling the origin; red line denotes migration trajectory in the new inputted embryo; misarranged cell is highlighted with green point, otherwise red.

**Figure S15.**
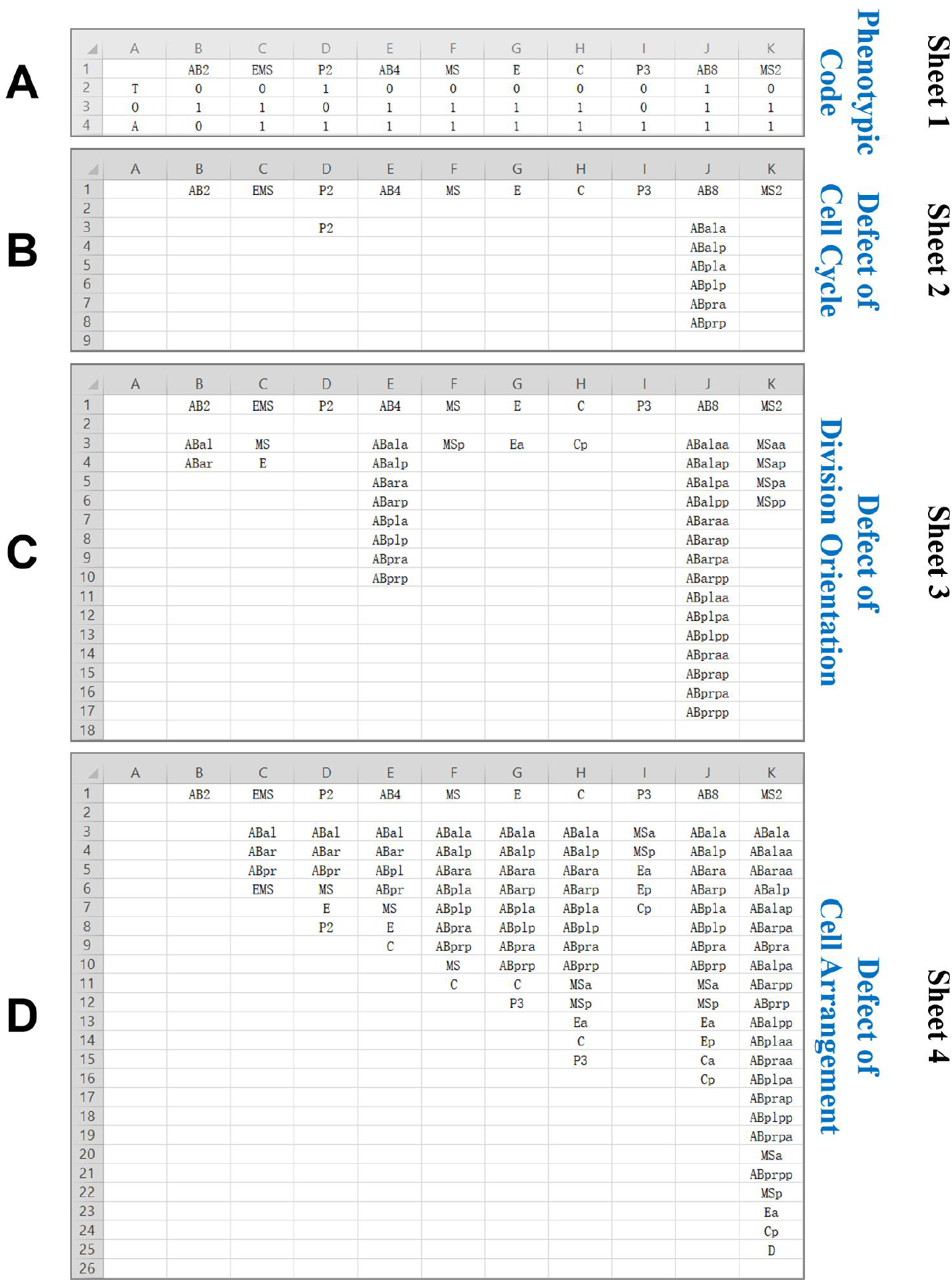
Reproducible defective phenotype of mutant from 4- to 24-cell stage, using *mex-5* as example. Each file consists of 4 sheets containing phenotypic code, defect of cell cycle, defect of division orientation and defect of cell arrangement respectively. The first row of each sheet is always the 10 division events listed successively according to their division timing and order. A. Phenotypic code formed by division timing *T_i_*, division orientation *O_i_* and cell arrangement *A_i_*. Each property would be assigned 1 if it’s defective, otherwise 0. B. Defect of cell cycle. Each cell listed is found to be defective in at least two mutant replicates of the gene. C. Defect of division orientation. Each cell listed is found to be defective in at least two mutant replicates of the gene. D. Defect of cell arrangement. Each cell listed is found to be defective in at least two mutant replicates of the gene.

**Figure S16.**
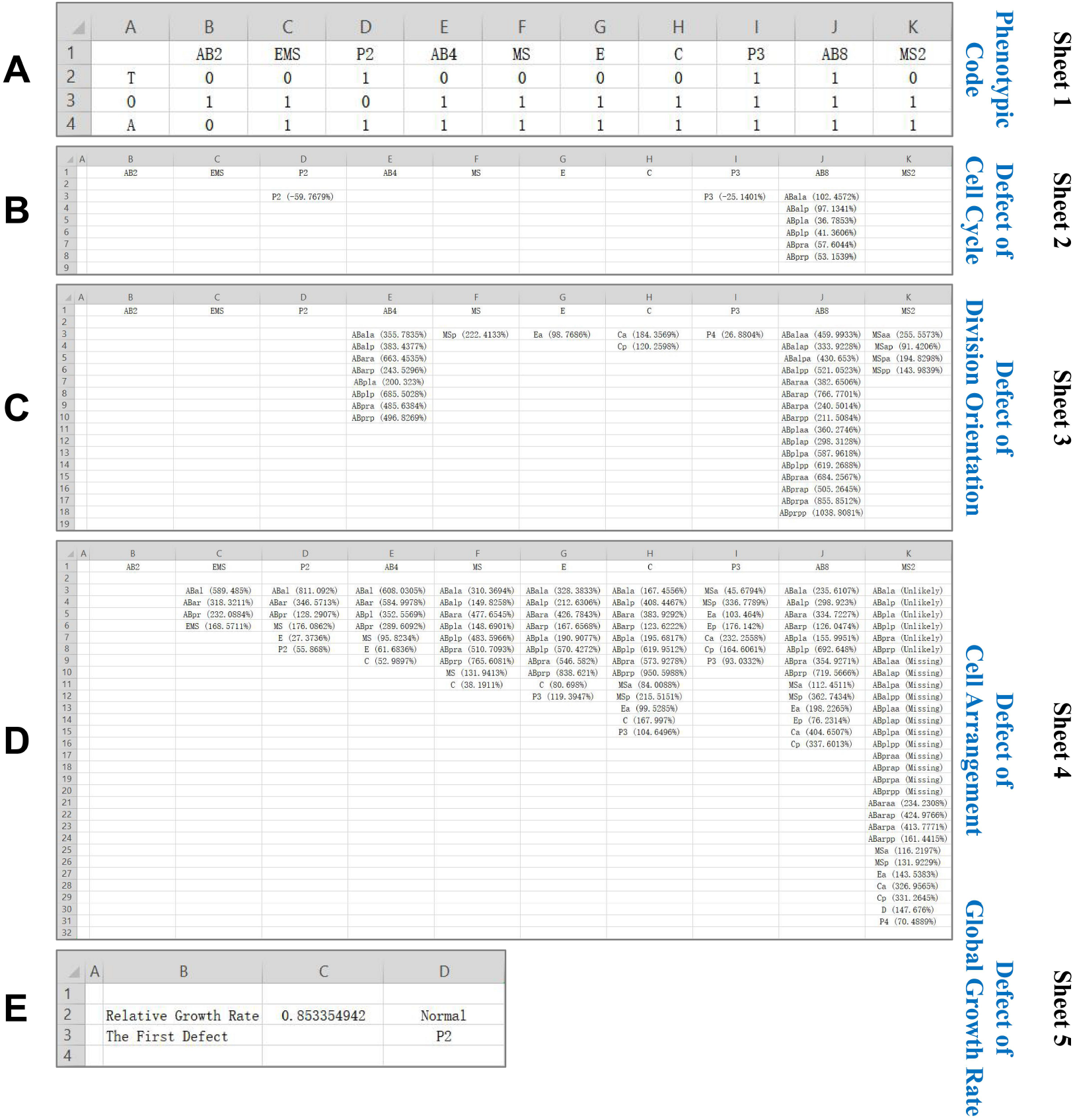
Quantitative phenotyping on defect of the 1818 mutant embryos from 4- to 24-cell stage, using *mex-5* as example (serial number 217, Table S6). Each file consists of 5 sheets containing phenotypic code, defect of cell cycle, defect of division orientation, defect of cell arrangement, and defect of global growth rate respectively. The first row of the first four sheets is always the 10 division events listed successively according to their division timing and order. A. Phenotypic code formed by division timing *T_i_*, division orientation *O_i_* and cell arrangement *A_i_*. Each property would be assigned 1 if it’s defective, otherwise 0. B. Defect of cell cycle. Each cell listed is found to have significantly longer or shorter cell cycle compared to the wild-type samples (99%), and the relative deviation 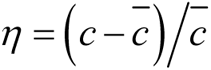 is also recorded. C. Defect of division orientation. Each cell listed is the daughter cells of the division events and found to be significantly defective compared to the wild-type samples (99%). The relative spatial deviation 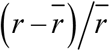 is also recorded (Fig.4F). D. Defect of cell arrangement. Each cell listed is found to be significantly defective compared to the wild-type samples (99%), and the relative spatial deviation 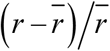 is also recorded. E. Defect of global growth rate. The relative growth rate is listed with its defective state (“Faster”, “Slower” and “Normal”, 99%) as well as the first division-timing defect (Fig.3B, Fig.S1D).

